# Neural representations supporting reuse of abstract structural knowledge in humans

**DOI:** 10.1101/2024.09.20.614119

**Authors:** Lennart Luettgau, Nan Chen, Tore Erdmann, Sebastijan Veselic, Zeb Kurth-Nelson, Rani Moran, Raymond J. Dolan

## Abstract

A human ability to adapt to the dynamics of novel environments relies on abstracting and generalizing from past experiences. Previous research focused on how humans generalize from isolated sequential processes, yet we know little about mechanisms that facilitate adaptation to more complex dynamics, including those that govern much everyday experience. Here, using a novel sequence learning task based on graph factorization, coupled with simultaneous magnetoencephalography (MEG) recordings, we ask whether reuse of experiential “building blocks” is reflected in neural representations that could support inference and generalization. Behavioral results were consistent with participants decomposing task experience into subprocesses, both abstracting their dynamical structure away from their sensory specifics and reusing these in a new task environment. Neurally, this was accompanied by a learning-induced, condition-specific, increase in neural similarity among stimuli conditional on their shared structural role in abstract subprocesses across distinct task phases. Consistent with this, a complementary dynamical role decoding analysis revealed enhanced neural decodability, across experiential contexts, for stimuli that adhered to identical structural transition types, specifically when prior experience included the relevant graph structure. Decoding strength for these role representations was positively related to behavioral success in reuse of subprocess knowledge, a relationship that will require future independent replication. The findings provide neural evidence consistent with the notion that abstract role representations may contribute to the reuse of experience within new contexts.

## Introduction

A defining feature of human cognition is an exceptional ability to adapt rapidly to novel contexts. An influential set of proposals suggests this relies on abstraction and generalization of past experiences (Allen et al., 2020; Dekker et al., 2022; Kumar et al., 2022; Lake et al., 2015, 2017; Lehnert et al., 2020; Tsividis et al., 2021) enabling application of experiential knowledge to entirely new situations (Behrens et al., 2018; Lake et al., 2017; Mark et al., 2020; Shepard, 1987). Thus, much research has focused on how humans detect (Turk-Browne et al., 2008), generalize (Reber, 1967) and neurally represent a single dynamical, and sequentially, unfolding process (Garvert et al., 2017; Henin et al., 2021; Schapiro et al., 2013; Sherman et al., 2020) or do so from piecemeal presentation of a graph structure (Rmus et al., 2022). Human neural evidence indicates these processes relate to representation of experiential elements in the hippocampal-entorhinal system (Garvert et al., 2023; Zheng et al., 2024). However, despite this progress, we know little regarding how humans abstract from the more complex dynamics that characterize much everyday experience.

Computational modeling with artificial neural networks, trained to perform multiple tasks (Johnston & Fusi, 2023; Yang et al., 2019), and cross-species neural evidence, propose a low dimensional neural coding of abstract, disentangled, task variables enables a linear readout of relevant task dimensions (Bernardi et al., 2020; Courellis et al., 2024). Brain structures implicated here include the hippocampal-entorhinal system (Baram et al., 2021; Bernardi et al., 2020; Courellis et al., 2024), frontal cortex (Baram et al., 2021; Bernardi et al., 2020; Courellis et al., 2024; Flesch et al., 2022, 2023; Zhou et al., 2021) and posterior parietal cortex (Flesch et al., 2022, 2023). Inference and generalization of knowledge could allow for a computationally efficient “recycling”, or reuse of learnt representations in novel contexts. Such a process would be akin to the abstraction of structural representations from sensory experiences that is proposed as implemented within the hippocampal-entorhinal system (Behrens et al., 2018; Whittington et al., 2020, 2022).

Previous studies have considered a single dynamical process/structure in relation to generalization to new situations. This ignores that our everyday world experience is often the product of complex environmental dynamics, comprising a *multitude* of simultaneously evolving structural subprocesses. Indeed, despite its ubiquity, we know little about the mechanisms of efficient adaptation to environmental dynamics generated from multiple simultaneous subprocesses (but see Conway & Christiansen, 2006 investigating behavior in response to mixed, but temporally well-separated stimulus sequences produced by two artificial grammars). Previously we found behavioral evidence for reuse of subprocesses that involved parsing complex task experience into component “building blocks”. Based on computational modeling and behavioral evidence, we proposed a cognitive-computational mechanism whereby structural mapping of prior experience facilitates reuse of knowledge in new tasks (Luettgau et al., 2024). While the latter data extended on previous behavioral findings regarding compositionality (Lake et al., 2015, 2017; Rubino et al., 2023), we lack a neural account of how this might be implemented.

In this study we hypothesized that adaptation to new experiences relies on reuse of abstracted representations of structural knowledge. Throughout this work, we use “generalization” to refer to the ability to apply learned relational structure to novel stimulus combinations within the same task context, and “reuse” to refer to the application of abstracted representations across distinct task contexts or learning phases (“transfer learning” denotes the second learning phase). In our task, reuse is most directly indexed by experience probes at transfer learning, which test knowledge of experienced transitions. Inference probes require reuse together with generalization to unobserved transitions within the transfer phase, and so provide a mixed measure. We use “abstract” to refer to representations that are invariant to the sensory specifics of individual stimuli, capturing relational structure (e.g., the role a stimulus occupies within a graph) rather than surface features. Our task has a compositional structure, with each experienced stimulus reflecting the combination of two simultaneously evolving subprocesses. We use ‘decomposition’ to refer to the process of parsing these compound experiences into their underlying components. We note that our study speaks most directly to decomposition and reuse of individual structural components, rather than to their recombination (or composition) into novel structures.

Under this framing, knowledge reuse involves mapping of inferred generative causes of an ongoing experience to support abstraction over concrete events (Mark et al., 2020; Pesnot Lerousseau & Summerfield, 2024; Whittington et al., 2020). We propose humans identify, neurally represent, and abstract the constitutive relational units of experience (“building blocks”). These representations may be reused to bias and support learning and inference about the relational units underlying new experiences, complementing rather than replacing direct experience. A representational reuse account of structural knowledge entails a prediction that, across different contexts, elements composing analogous subprocesses would show representational similarity, indicative of a shared abstract representation that could support reuse of experiential “building blocks”.

Prior studies have mapped the spatial location of low-dimensional neural representations of abstract and disentangled task variables to brain areas that include the hippocampal-entorhinal system (Baram et al., 2021; Bernardi et al., 2020; Courellis et al., 2024; Nieh et al., 2021), frontal cortex (Baram et al., 2021; Bernardi et al., 2020; Courellis et al., 2024; Flesch et al., 2022, 2023; Zhou et al., 2021) and posterior parietal cortex (Flesch et al., 2022, 2023). In parallel, M/EEG studies reveal that abstract, format-independent representations can emerge dynamically over time. For example, multivariate analyses of MEG data demonstrate cross-decoding of numerical magnitude across distinct symbolic formats (e.g., digits and dice), indicating a shared, format-invariant representation that unfolds with distinct temporal dynamics (Teichmann et al., 2018). More broadly, M/EEG evidence suggests that task-relevant information can be dynamically prioritized, compressed, and reformatted, indicative of a rapid temporal evolution of abstract and behaviorally relevant representations even when spatial localization remains coarse. Recent MEG data suggests that high-dimensional sensory inputs are progressively transformed into lower-dimensional, task-relevant, feature manifolds across distinct processing stages, reflecting dynamic gating and reformatting of information in support of flexible categorization (Duan et al., 2024). Complementary EEG–fMRI evidence also indicate abstract task goals are encoded in a low-dimensional neural geometry that emerges in frontal cortex and is communicated to posterior regions to guide implementation of stimulus-specific representations, with the fidelity of this geometry predicting behavioral performance (Zhang & Yu, 2024). Likewise, converging behavioral and computational data indicates successful reuse and generalization depends on detecting and encoding abstract relational structure rather than stimulus surface features. In effect, training enables transfer when task phases share an underlying rule or abstract relationship, even across modalities, but not when an overlap is purely perceptual (Garner et al., 2016).

At the same time, reuse of shared abstract representations introduces a computational trade-off, such that overlapping codes that facilitate transfer can also induce interference, as observed in both humans and artificial neural networks (Holton et al., 2025). However, while there is now much work elucidating how learned representations are generalized, it remains unclear whether and how abstracted structural representations are expressed neurally during ongoing stimulus processing, and how they are shaped by prior structural experience. Thus, beyond characterizing where abstract representations are located in the brain there is a related question of how they dynamically unfold in support of reuse of structural knowledge across contexts.

To address these questions, we developed a sequence learning paradigm based on the graph theoretical concept of graph factorization, emulating environments whose dynamics are the product of multiple simultaneous subprocesses. Product graphs (Cartesian product), wherein the joint dynamics of two or more graph factors create complex product state space dynamics that can be factorized (and simplified) into constitutive factors, provides a compelling and flexible test bed for probing structural knowledge reuse of multiple (independent) and simultaneously unfolding subprocesses. Examples of such graph products and their factorizations, from a vast space of possible combinations, are provided in Figure 1A. Here we take inspiration from prior studies in neuroscience (e.g., Barron et al., 2013; Rubino et al., 2023; Schwartenbeck et al., 2023) and a prior cognitive literature, e.g., recursive operations over hierarchies (O’Donnell et al., 2009, 2011), hierarchical function composition, or rich forms of semantic composition in language to create complex concepts (Murphy, 1988).

**Figure 1.**
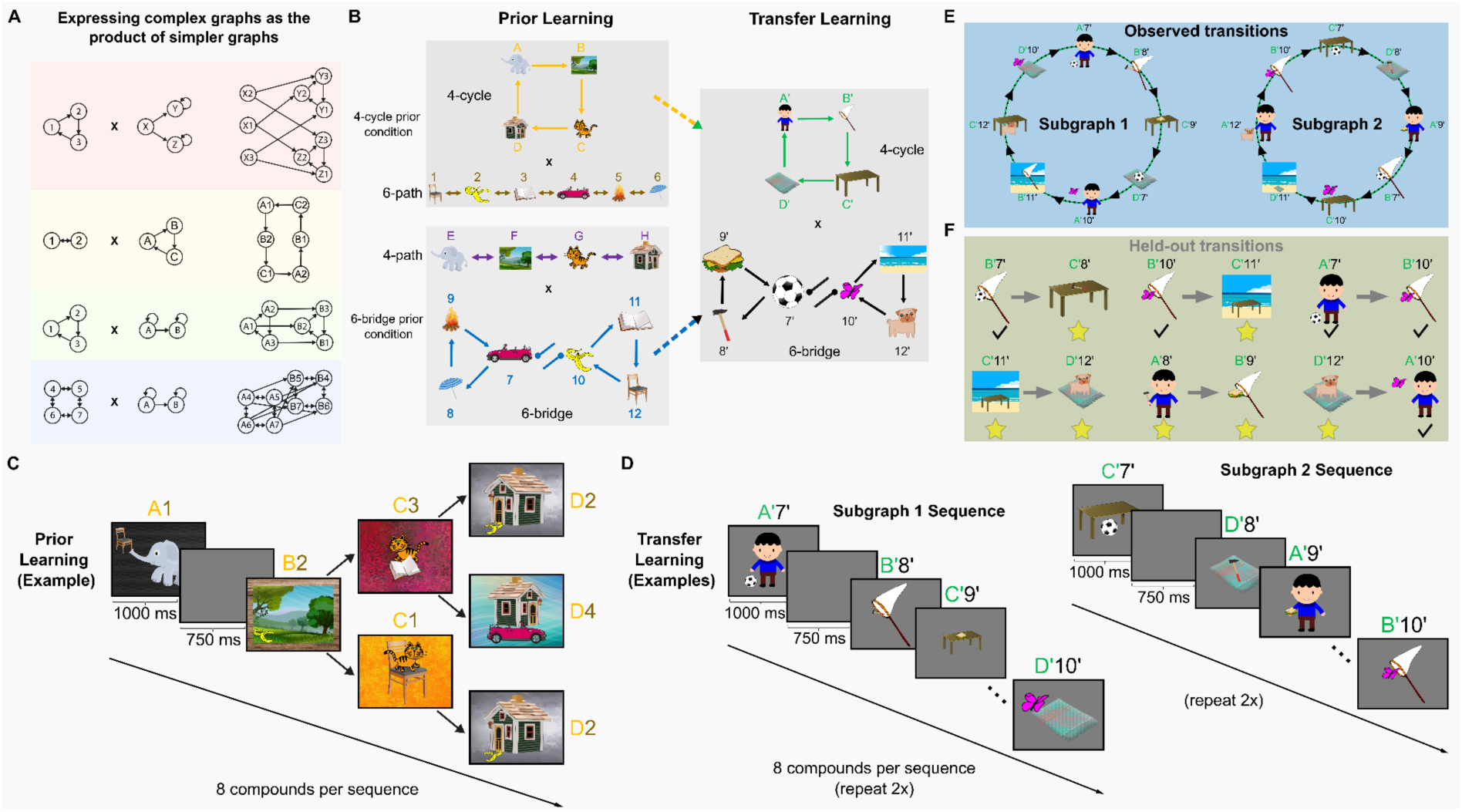
Experimental design. **A)** Exemplar product graphs (Cartesian product, right) selected from an array of possible graph products that can be constructed from two simpler graph components (left). The graph product of two or more simple graph components results in complex product state spaces, which can be decomposed (and simplified) into their constituting components. **B)** Graph factors used in prior learning (left) and transfer learning (right) **C)** During the sequence learning task (both prior and transfer learning), graph factors were combined into compound graphs (product graphs) by merging their images. In both prior learning and transfer learning, participants were presented with sequences of meaningful compound holistic images, such as a cat on a chair or a house with a car. Depicted is an example sequence generated by the product graph of the yellow graph factor and the gold graph factor used during prior learning (panel B, left, top). The example sequence involved a simultaneous traversal in the yellow graph factor and in the gold graph factor, as shown in panel B (left, top). After each sequence presentation (consisting of 8 compound stimuli), participants’ knowledge of the graph structures was assessed in two ways, as detailed in Figure 2. **D)** During sequence presentations in transfer learning, participants only observed a subset of transitions from the compound graph formed by the green graph factor and the black graph factor, as shown in panel B (right), e.g., the 16 depicted transitions, forming two disjoint subgraphs. The remaining possible compound graph transitions formed a held-out set (see examples in panel F). This allowed us to test participants’ understanding of the subprocesses generating the observed sequences. The same principles applied to prior learning. Depicted are example sequences produced by the subgraphs of the product graph used in transfer learning, involving a simultaneous traversal in the green graph factor and in the black graph factor, as shown in panel B (right). Subgraph sequence 1 (left) was presented two times, after which subgraph sequence 2 (right) was presented twice. **E)** For illustration of the transfer learning compound graph, we only show the parts of the compound graph for transfer learning that participants experienced during sequence presentation, created from the 4-cycle and 6-bridge graphs shown in panel B (right). Note that the 6-bridge graph did not feature a truly bi-directional edge between the central nodes (7//7’ and 10/10’), but instead had a “refractory” edge, such that it was not possible to transition back and forth within one step between central nodes of the graph. In both phases of the task (prior and transfer learning), the compound graphs consisted of 24 compound images. **F)** Examples of held-out transitions between compound images used in the transfer learning phase. Note these entailed transitions between two compounds which were observed (as part of other transitions) during sequence presentations (denoted by a black tick below), or between an observed and a compound that had never been presented (denoted by yellow stars below) or between two new compounds. In any case, the specific transitions between compounds had never been observed during sequence learning.

Using magnetoencephalography (MEG) we ask three specific research questions. First, do stimuli that share the same abstract subprocess become more similar neurally following learning, and is this representational change specific to the subprocess experienced during prior learning? Second, do neural representations also come to reflect the fine-grained dynamical roles stimuli occupy within a subprocess? Third, does the strength of these emergent neural representations relate to participants’ ability to successfully reuse subprocess knowledge in new contexts? Addressing these questions allowed us to characterize the neural representational basis of reuse of structural knowledge, establishing when during stimulus processing abstract structural knowledge is expressed and how it is shaped by prior experience. We found that distinct sensory stimuli showed greater neural similarity when they adhered to the same underlying abstract relational dynamics, while distinct stimuli in analogous “dynamical roles” were represented similarly across entirely different task phases. Decoding strength for these dynamical roles was positively related to performance on experience probes for the re-experienced structural component, but not on inference probes, and this relationship did not differ significantly from that for the other component.

## Results

In brief, participants (*N* = 50, *N* = 25 subjects per condition) performed a sequence learning task, observing image sequences produced by walks on product graphs constructed from smaller graph factors (Fig. 1A and B). The experiment consisted of two phases (prior learning and transfer learning, see task design details in Figure 1 and 2). During both task phases, participants viewed sequences of compound images comprising two elements, depicting meaningful scenes. The compound images used for the sequences were generated from combining images on two different graph factors (Fig. 1B), where these differed between-subjects and across the two task phases.

**Figure 2.**
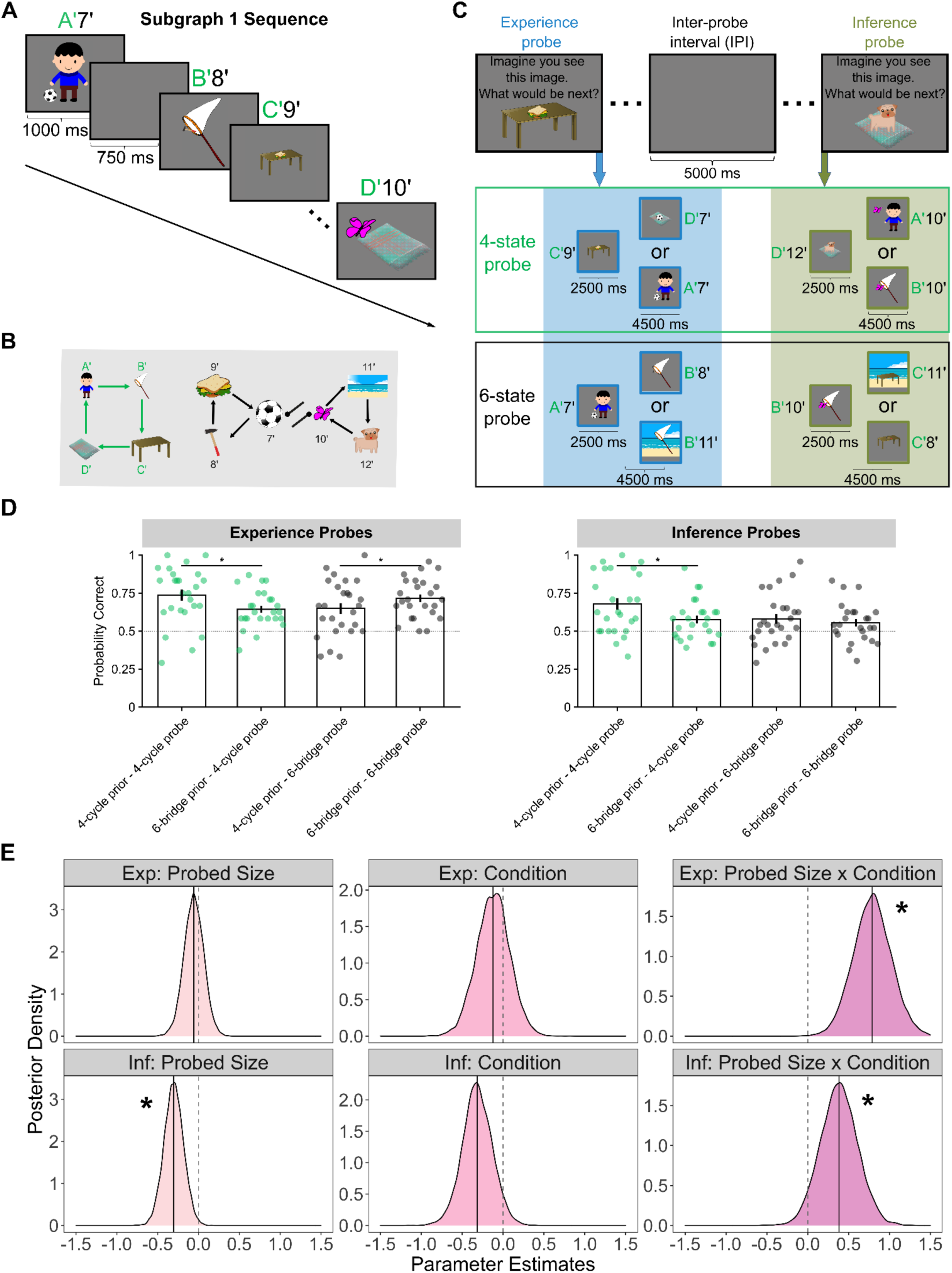
Behavioral task and transfer learning performance. A) Example sequence produced by subgraph 1 of the product graph used in transfer learning (reproduced Fig. 1D left). B) The compound image sequence in panel A was generated from a simultaneous traversal of two *graph factors,* by combining their images (reproducing Fig. 1F for convenience). The example sequence was generated from a 4-state cyclic graph factor and a 6-state bridge graph factor (reproducing Fig. 1B for convenience). C) Exemplar schematic of experience and inference probes during transfer learning, respectively. During both prior and transfer learning, following presentation of an entire compound image sequence, participants were probed to predict upcoming states that adhered to one of the 16 experienced compound image transitions (experience probes, left column). Additionally, an inference probe (right column) tested participants’ ability to infer and accurately predict held-out transitions (Fig. 1C). In both probe questions, participants saw a compound image and were asked: “Imagine you see this image. What would be the next image?”. Two compound images – correct next compound image (eligible based on the graph structure) and a lure compound image (non-eligible) – were presented as choice options. The lure compound image always matched the correct option on one of the two individual images composing the compound image (e.g., D’12’ -> A’10’ or B’10’). This allowed us to test knowledge of both graph factors separately (in the current example, knowledge that D’ allows a transition to A’ but not B’) (top row: 4-state factor; bottom row: 6-state factor) underlying the observed sequence (and the held-out transitions) separately. D) Aggregated probabilities of correct answers during the transfer learning phase, separately for experience (left) and inference probes (right) as a function of probed size (4-state cycle (green) vs 6-state bridge (black)) and condition (Condition 4 vs Condition 6). Each dot represents the arithmetic mean of experience and inference probe performance for one participant, bars represent the arithmetic mean of the distribution, error bars depict the standard error of the mean and the dashed gray line represents chance level point estimate (probability correct = 0.5). Asterisks indicate condition differences for which contrasts of posterior parameter estimates exclude 0. E) Posterior density plots for each parameter estimate (posterior mean = black line) for the best-fitting GLM. The dashed vertical line represents a zero effect. In probed size (Component 4 or Component 6) and probe type (experience vs inference) effects, parameter estimates below zero indicate higher accuracy in Component 4 probes (vs. Component 6 probes) and experience probes (vs. inference probes), respectively. In the condition effect, parameter estimates below zero indicate higher accuracy in Condition 4 (vs. Condition 6). The positive interaction effect indicates selectively higher accuracy in Component 4 probes in Condition 4, and higher accuracy in Component 6 probes in Condition 6. Asterisks indicate GLM parameter estimates for which posterior distributions exclude 0.

Using a between-subjects design, in a prior learning phase, participants in a 4-cycle condition experienced sequences generated from the product of a 4-state cycle graph factor and a 6-state path-graph factor (Fig. 1B top panel, left, Fig. 1C for example sequences). Likewise, participants in a 6-bridge prior condition saw compound image sequences produced by combining a 4-state path-graph factor and a 6-state bridge graph factor (Fig. 1B bottom panel, left). Each of these graph factors is considered a structural “building block” underlying participants’ holistic experience.

The prior learning phase was followed by a transfer learning phase, identical for both conditions (Fig. 1D), but now compound image sequences were composed from entirely novel picture elements. The latter were the product of a 4-state cyclic graph factor and a 6-state bridge graph factor, respectively (Fig. 1B right). This between group design meant that, for each participant, the structure of only one of the two graph factors (“building blocks”) that generated the compound image sequences was common across the prior and transfer learning phase (e.g., in the 4-state cycle prior condition, the 4-state cycle, but not the 6-state bridge, was present in both prior and transfer learning, Fig. 1B). Importantly, these graph factors were embedded by entirely different images across both task phases, rendering any learning unlikely based on stimulus similarity.

For brevity and clarity, we hereafter refer to the two prior learning conditions as Condition 4 (4-cycle prior) and Condition 6 (6-bridge prior), based on which graph structure each contributes to the transfer learning phase as a recurring structural component. Correspondingly, we refer to the 4-state graph factor (i.e., the 4-cycle) as Component 4 and the 6-state graph factor (i.e., the 6-state bridge) as Component 6. Under this notation, Condition 4 participants can reuse Component 4 knowledge during transfer, while Condition 6 participants can reuse Component 6 knowledge.

Our central prediction was that individuals would abstract experienced subprocesses (i.e., graph factors) from prior learning and reuse this knowledge during transfer learning, benefiting from recurrence of a previously experienced subprocess. To test this, in both task phases (i.e., prior and transfer learning), participants experienced sequences of a subset of all possible transitions between compound images (Fig. 1E and F illustrate this as an example for the transfer learning phase product graph). The remaining possible transitions between compound images (based on transitions on the underlying graph components) formed a held-out set of transitions between compound images (see examples in Fig. 1D). Leveraging held-out transitions enabled us to test knowledge for unobserved transitions between compound stimuli, at each task phase (i.e., prior and transfer learning). In essence this provided an assay of whether participants had acquired a factorized representation, i.e., could generalize to unobserved transitions (knowledge of the individual graph factors producing the experienced compound sequence (Fig. 1E/F)). For details on the transitions, we refer to the Methods section.

During prior and transfer learning, “experience probes” tasked participants to predict upcoming states that followed one of the compound images from the experienced subgraph (Fig. 2C, left column). Likewise, in both prior and transfer learning “inference probes” (Fig. 2C, right column) tested for an ability to generalize to held-out transitions, i.e., to make accurate predictions about transitions never directly experienced. In both experience and inference probes, participants were tasked to choose between the correct next compound image (correct transition based on the generative graph structure) and a lure (incorrect transition) compound image. The lure compound image always matched the correct compound image with respect to one of the component individual images (e.g., D’12’ -> A’10’ or B’10’), allowing us to test knowledge pertaining to each of the graph factors separately (here, the knowledge that in the 4-state graph factor component D’ is followed by A’ rather than B’). Probe questions were carefully designed in such a way that did not allow to infer the structure from these questions (see Methods for details). Thus, the task allowed us to test if subjects utilize the structural “building blocks” underlying their experience (graph factors, subprocesses). Additionally, we could test whether knowledge of subprocess dynamics is reused in novel contexts. This is because in our design, despite using entirely different stimulus sets across the task phases, one of the abstract graph factors was consistent for participants in both conditions, allowing reuse of this specific structural knowledge. However, knowledge about the other graph factor that generated experiences in prior learning was irrelevant, enabling control for general learning effects.

In summary, participants viewed sequences of images that were combined from concurrent walks on two underlying graph structures (“building block”). In prior learning, they learned image sequences generated from two graph structures. In a subsequent transfer learning phase, participants saw new image sequences, but one of the two structural “building blocks” that underlie the observed image dynamics remained the same for each participant. Throughout both phases, participants were asked to predict upcoming images and infer never observed transitions. This task feature allowed us to test reuse of prior structural knowledge, which we predicted would reflect in higher accuracy in the consistently encountered “building block” (but not the other, irrelevant “building block”) – providing evidence for a hypothesis that participants reuse components of learned structural dynamics despite changes in component sensory elements.

### Abstracting and reusing subprocesses

A behavioral prediction was that individuals abstract experienced subprocesses (i.e., graph factors) from prior learning and reuse this knowledge during transfer learning. Thus, at transfer learning Condition 4 participants should perform better on Component 4 probes compared to Condition 6 participants, and vice versa for Component 6 probes. This predicts at transfer learning a disordinal interaction where the probability of being correct varies as a function of prior learning condition (Condition 4 vs Condition 6) and probed size (Component 4 vs Component 6) (Fig. 2D). To model candidate processes that might generate the observed transfer learning behavioral data, we defined four Bayesian multilevel GLMs (GLM1–4, Eq. 1-4). The most parsimonious model (GLM4, see Methods for model comparison) indicated the effects for probed size and prior learning condition on accuracy were qualified by a three-way interaction effect between these two variables and probe type (experience vs inference). In post-hoc GLMs, we observed non-zero interaction effects between probed size and condition (experience probes: µ_SIZExCOND_ = .79, credible interval (CI) = [.42; 1.17], Fig. 2E top row, third panel; inference probes: µ_SIZExCOND_ = .38, CI = [.02; .75], Fig. 2E bottom row, third panel)

Disentangling these effects using post-hoc contrasts [Condition 4 – Condition 6], showed that experience probes performance was higher for Component 4 probes in Condition 4 (vs Condition 6; Fig. 2D left panel, green dots; mean difference of 10,000 posterior samples: .36, CI = [.17; .56]) whereas for Component 6 probes, performance was superior in Condition 6 (vs Condition 4; Fig. 2D left panel, black dots; mean difference of 10,000 posterior samples: -.23, CI = [-.42; -.04]). In inference probes, performance on Component 4 probes was higher in Condition 4 (vs Condition 6; Fig. 2D right panel, green dots; mean difference of 10,000 posterior samples: .38, CI = [.20; .56]). For Component 6 probes, there was no evidence for a performance difference between conditions (Condition 4 – Condition 6; Fig. 2D right panel, black dots; mean difference of 10,000 posterior samples: -.11, CI = [-.26; .04]). Posterior distributions for the other parameter estimates of these GLMs are depicted in Figure 2E. Splitting behavioral results by performance levels – high performers (>50th percentile of overall prior and transfer learning performance distribution) vs low performers (<=50th percentile) – yielded similar results (Supplementary Fig. 1).

These findings replicate previous results from a larger cohort using a similar design (Luettgau et al., 2024). The Component 6 experience probe behavior utilized here indicates subjects not only extract and reuse subprocess dynamics for cyclical structures (as we previously showed), but also for more complex dynamics (bridge graphs). The observed pattern is not accounted for by general practice or non-specific meta-learning related improvement effects (as such accounts would predict similar performance for both probed sizes across conditions). Instead, the results are consistent with participants being influenced by experiential subprocess dynamics (Luettgau et al., 2024), where generative consistency across both task phases biases performance during transfer learning, though direct experience remains a dominant driver of behavior

### Structural abstraction and reuse of subprocesses

A representational reuse account predicts elements of analogous subprocesses, across different experiential contexts, would show a representational similarity reflective of adherence to a shared abstract representation. Thus, this analysis addresses our primary research question as to whether abstract structural representations are expressed neurally during stimulus processing, and whether their emergence is shaped by prior learning experience. To test the above hypothesis we recorded MEG signals during distinct localizer tasks, from before and after sequence learning (PRE and POST localizer task, Fig. 3A). We reasoned that a representational similarity in experiential “building blocks” should be reflected in a task-induced (PRE to POST) increase in neural similarity between distinct stimuli as a function of adherence to the same (compared to different) underlying relational structure, in this case adhering to identical subprocess dynamics (Fig. 3B). To test this, we used a feature similarity analysis (akin to representational similarity analysis (RSA) (Kriegeskorte et al., 2008) on MEG data (*N* = 50, one subject from the behavioral sample was excluded due to MEG artifacts) from PRE and POST localizer task sessions (Fig. 3A).

**Figure 3.**
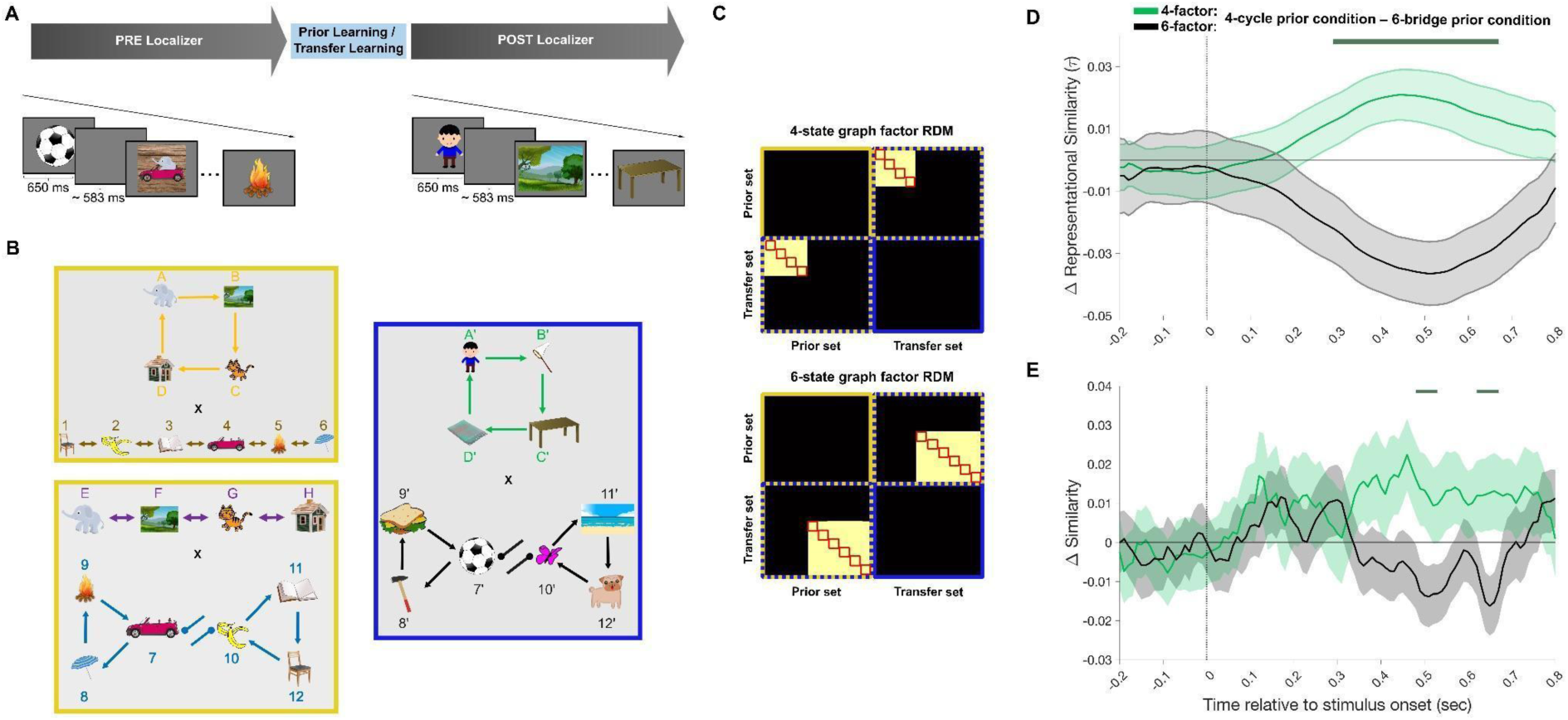
Neural representation of structural mapping between subprocesses. A) Participants performed two stimulus localizer tasks, once before and one after (PRE and POST localizer) the main experimental task (prior and transfer learning). These tasks comprised multiple presentations of all features used in prior and transfer learning (PRE and POST). Additionally, the PRE localizer also contained the compound images used during prior learning (not relevant for this analysis). A feature similarity analysis (akin to RSA) on stimulus-evoked neural activity recorded during the two localizer tasks (PRE and POST localizer) for the two feature sets, comprising the compound images used in prior and transfer learning at time-points from -200 to 800 ms relative to stimulus-onset. B) Feature sets in prior learning phase (yellow frame, left) and transfer learning phase (blue frame, right). C) The presented theoretical representational dissimilarity matrices (RDM) were regressed onto the empirical neural similarity matrices – separately for the 4-state graph factor features and the 6-state graph factor features. Lighter colors denote higher values. Note that the hypothesized similarity changes are computed across the two feature sets used in prior learning (yellow) and transfer learning (blue), but not within each set, avoiding confounding task-related changes with feature similarity unrelated to abstract subprocess structure. Red squares indicate structural mappings between stimulus sets (e.g., stimulus in the first, second, third position in the 4-state cycle), applied in E). D) Time courses of condition differences (Condition 4 – Condition 6) on Component 4 (green) and Component 6 (black) feature similarity changes in the weighted average PCs from PRE to POST localizer. Shaded error bars indicate standard errors of difference. Dark green line represents time points at which feature similarity changes for the condition x graph factor interaction effect were statistically significant, corrected for multiple comparisons at the cluster-level (P_FWE_ < 0.05, non-parametric permutation test). For prior learning group and graph factor main effects all *P_FWE_* > 0.118 (non-parametric permutation test). Horizontal line indicates a null effect. E) Time courses of condition differences (Condition 4 – Condition 6) in structural role–based representational similarity on Component 4 (green) and Component 6 (black), computed from sensor-level RSA in the localizer task (POST – PRE). Structural role similarity reflects increased neural similarity for stimuli occupying the same graph-theoretic role (e.g., boundary, intermediate, central) within each graph structure. Shaded error bars indicate standard errors of the difference. The dark green line denotes time points at which the prior condition × graph factor interaction was statistically significant, corrected for multiple comparisons at the cluster level (P_FWE_ < 0.05, non-parametric permutation test). For prior learning group and graph factor main effects all P_FWE_ > 0.90 (non-parametric permutation test). The horizontal line indicates a null effect.

As an initial step, to denoise sensor-level neural signals in a principled manner, we implemented a dimensionality reduction approach (principal components analysis, PCA, see Methods for details). Thus, for each participant, we implemented the following steps separately for PRE and POST localizer sessions. First, we extracted the minimal number of PCs that accounted for >=80% of the variance of neural activity across sensors for all trials and time points (-200 to 800 ms relative to stimulus onset). Next, for each time point, trial and PC we calculated the neural PC activity (i.e., PC score). By averaging PC activity across all trials presenting the same feature (i.e., element images of graph factors) we obtained an activity index for each PC, feature and time point. We next computed the dissimilarity between each pair of features as the absolute difference in their activity indices. This provided for a Representational Dissimilarity Matrix (RDM) at each time point, and for each PC. Next, for each time point, we calculated an across PC weighted average RDM (with weights being variance explained by each PC relative to the total variance explained by all PCs included in the analysis), resulting in a weighted average RDM for each participant, time point and session (PRE/POST). We then subtracted the POST weighted average RDM from the PRE weighted average RDM for each participant to obtain a PRE-POST difference weighted average RDM. Finally, to quantify feature similarity change at each time point, and for each participant, we computed Kendall’s tau correlation coefficient between theoretical RDMs and PRE-POST differences in neural RDMs (similar to Luyckx et al., 2019; Nili et al., 2014). Our theoretical RDMs relate to a hypothesized abstract representation of subprocesses that reflect higher similarity between feature-pairs, where features belong to different learning phases (one from primary, the other from transfer) but share a structural graph component (separate design matrices for the 4-state graph factor and 6-state graph factor, Fig. 3C).

Within the PRE-POST change in weighted averaged RDMs, across participants we observed a qualitative pattern indicative of increased neural similarity between stimuli that shared the same underlying subprocess across task phases. A group-level GLM (including condition, graph factor main effects and a condition x graph factor effect) revealed that highly distinct stimuli that adhered to the same abstract relational structure (4-state cycle and 6-state bridge graphs, respectively), manifest a significantly greater neural similarity (Fig. 3D). For illustration purposes, we computed the average difference between feature similarity metrics for both prior learning conditions (4-cycle prior condition – 6-bridge prior condition), separately for each of the graph factors (see Fig. 3D). There was a statistically significant interaction effect of condition x graph factor spanning approximately 300 – 680 ms post-stimulus onset (cluster-level *P*_FWE_ = .004, corrected for multiple comparisons using non-parametric permutation tests, see Methods for details). The timing of this effect indicates that abstract structural information is expressed relatively early during stimulus processing, within a window broadly consistent with higher-order representational processing rather than early sensory encoding or late decision-related activity (Nour et al., 2021). Importantly, this time window should be interpreted as a marker of when abstract structural representations are neurally available, not as evidence for a discrete computational stage. We did not observe prior learning group and graph factor main effects (all *P_FWE_*> 0.118, non-parametric permutation test). Post-hoc tests of condition differences for each graph factor indicated that within the time period where we observed a significant condition x graph factor effect, participants in Condition 4 showed higher positive change in feature similarity that mapped to Component 4 across both task phases, by comparison to participants in the Condition 6 (who experienced a 4-state path-graph in prior learning, Fig. 3D, green line, cluster-level *P*_FWE_= .040, non-parametric permutation test), and vice versa for the Condition 6 (Fig. 3D, black line; dark green line in Fig. 3D represents time points at which feature similarity changes for both post-hoc contrasts are significant; cluster-level *P*_FWE_ = .002, non-parametric permutation test). Performing the former analysis with an unweighted average across the PCs that accounted for >= 80% of the variance (all PCs contribute equally) yielded highly similar results (interaction effect of condition x graph factor from 320 – 690 ms post-stimulus onset; cluster-level *P*_FWE_ = .007, corrected for multiple comparisons using non-parametric permutation tests). Importantly, to verify that the chosen PCA approach did not impose artificial structure within the data, or introduce other potential confounds, we also conducted the above analyses on sensor level MEG data. Highly similar results were obtained, with consistent results for different similarity metrics (absolute distance (L1 norm) and Euclidean distance (L2 norm), Supplementary Figure 3)), demonstrating that our results and conclusions are robust across analytic choices.

These results are not explained by participants simply learning to differentiate which of the 10 observed features pertain to Component 4 and 6, since this predicts only a main effect of graph factor on neural similarity (features within one graph factor being represented more similarly than across graph factors), but not an interaction effect of condition x graph factor.

The observed increase in neural similarity between stimuli associated with the same graph factor could, in principle, reflect a more general categorical grouping of stimuli belonging to the same graph factor, rather than reflecting an abstract graph representation. To address this alternative account, we examined whether neural similarity in the localizer task is specifically structured according to the graph-theoretic role each stimulus occupies within its graph, using a finer-grained theoretical RDM that distinguishes between boundary, intermediate, and central positions. Structural roles were derived from each participant’s learning task sequences and defined by graph topology. For Component 6, roles were assigned using a bridge-graph traversal template (for Condition 6, whose prior learning involved the bridge graph) or degree-based assignment (for Condition 4, whose prior learning involved a line graph): boundary nodes were defined as nodes with degree 1 (graph periphery); intermediate nodes as neighbours of boundary nodes; and central nodes as the remaining, more densely connected nodes. For Component 4, two roles were distinguished, boundary-like nodes (those encountered first and last in the sequence traversal) and central-like nodes (those encountered second and third), reflecting structural position within the graph rather than arbitrary temporal sequence. For further details of role assignment, see Methods. This analysis revealed a cluster-based permutation-corrected interaction (*P_FWE_* < 0.05, non-parametric permutation test, Figure 3E) between prior learning group and graph factor at overlapping time windows with the previously reported PCA and sensor-level analyses investigating feature-mappings to abstract graph factors. We did not observe significant and multiple comparisons corrected prior learning group and graph factor main effects (all *P_FWE_* > 0.90, non-parametric permutation test). These results indicate that representational similarity increased selectively for stimuli sharing the same structural role in the graph structure relevant to each group’s prior experience. Critically, this effect was not observed uniformly for all stimuli associated with the same graph, arguing against a purely categorical grouping explanation. Instead, these findings provide evidence that learning induces representations that reflect higher-order relational structure, capturing graph-theoretic roles that are invariant across surface features of the embedding stimuli and are consistent with reuse across learning contexts.

Taken together, our findings indicate a task-induced, learning-related increase in neural similarity for stimuli sharing a structural role in subprocesses common across task phases and is compatible with a role representation reuse account, though we do not claim these representations are fully context-independent.

### Abstract representation of dynamical roles in subprocesses

The preceding analyses showed that learning increases neural similarity between stimuli that share a subprocess across task phases, with this increase being specific to stimuli occupying the same structural role (boundary, intermediate or central) within the graph rather than a uniform similarity increase across all stimuli belonging to that graph, arguing against a purely categorical account. However, those analyses index static stimulus representations measured passively, before and after learning. Here we ask whether role information is also expressed during the task itself, as participants process sequences, and whether information acquired during prior learning is reused in transfer learning. Specifically, we test whether neural activity following a transition carries information about the dynamical role of that transition. That is, whether the signal carries information about its position within the transition structure of the graph, such that a classifier would generalize to transitions involving entirely different stimuli (but sharing the same transition role) during transfer learning. If roles are reused across contexts, this generalization should be stronger in participants whose prior learning involved the relevant graph structure. Note that “generalization” in this section is used in the machine-learning sense: a classifier trained on neural activity evoked by one set of stimuli (prior learning) is tested on activity evoked by entirely different stimuli (transfer learning). This describes the neural signal, namely whether it carries role information that is shared across stimulus sets, and differs from the cognitive sense of generalization defined in the Introduction. We hypothesized that stronger cross-stimulus decoding in participants whose prior learning involved the relevant graph structure is neural evidence consistent with reuse. This analysis also provides the neural measure that allows us to relate role representations to behavior (see next section). Note that this can only be ascertained for Component 6, as this graph alone incorporates transitions of categorically distinct types.

To test this, we focused on MEG data recorded during prior and transfer learning phases involving, in Component 6, compound stimulus presentations that adhered to specific abstract transition types (Figure 4A). Within Component 6, these transitions comprise three distinct types, reflective of their graph locations namely transitions between boundary and intermediate graph nodes (1), transitions between intermediate and central graph nodes (2) and transitions between central and central graph nodes (3) respectively (see Methods for details). During prior learning, we trained classifiers on neural activity from stimuli belonging to each of these three transition types. In the transfer learning phase, we tested this generalization of pre-trained classifiers to identical transition types, but where these transitions now entailed entirely different stimuli. We used two-sample t-tests to ascertain classification accuracy between participant groups who had experienced different graphs during prior learning (Condition 4 experiencing a 6-path graph and Condition 6 experiencing a 6-bridge graph). This included multiple comparison correction based on nonparametric permutation tests.

**Figure 4.**
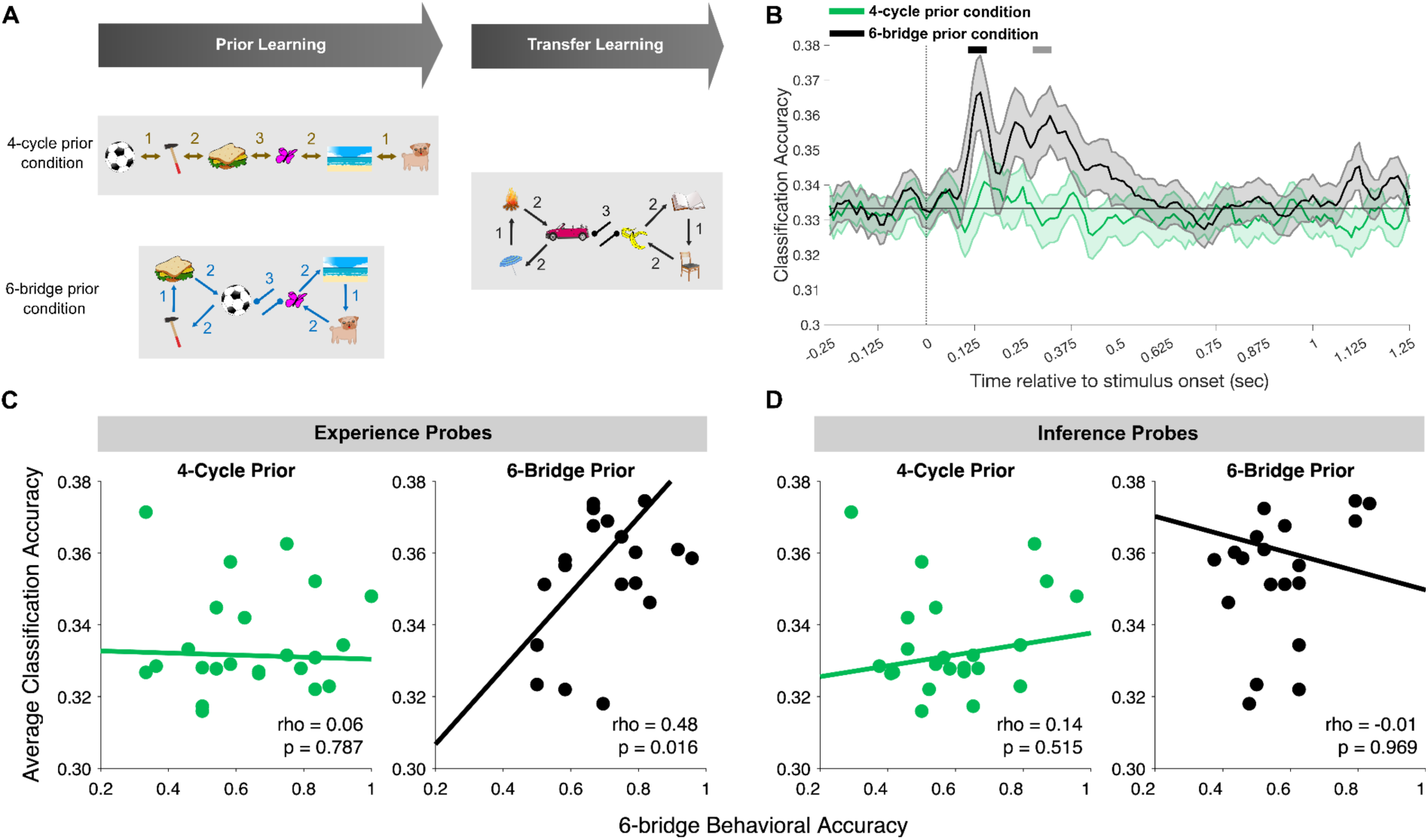
Abstract representation of subprocess dynamics. A) Knowledge reuse of decomposed subprocesses, discovered during prior learning, is related to neural similarities when subprocess dynamics are shared across prior and transfer learning. During prior learning, we trained multivariate classifiers (L1-regularized logistic regressions) to distinguish between different transition types in the 6-state graph factor for 4-cycle prior condition (6-path graph, top) and 6-bridge prior condition (6-bridge graph, bottom). We then tested the classifiers’ generalization ability on entirely new stimuli during transfer learning. Each number denotes a different type of transition, reflecting edges connecting graph nodes, namely boundary (1), intermediate (2), and central (3). During prior learning, classifiers were trained at post-transition stimulus presentations (e.g., in the 6-bridge prior, the classifier for transition type 2 was trained on neural activity recorded during presentations of the ball if preceded by the sandwich; or on the hammer if preceded by the ball). At transfer learning we tested the classifier’s generalization performance for identical transitions, but where this now involved entirely different stimulus sets. Since training and testing were performed on independent datasets this obviates cross-validation. Note that a comparison of classification accuracy across conditions was only possible for the 6-state graph factor, as the 4-state cycle does not provide for a distinction between transition types. B) Transfer learning time courses of 6-state graph factor transitions classification accuracy, averaged per condition based on prior experience of either 4-cycle (green) or 6-bridge condition (black). Shaded error bars indicate standard errors of the mean. Black and grey bars above indicate temporal clusters in which classification accuracy differences between conditions (6-bridge prior condition > 4-cycle prior condition) are significantly different (two-sample t-test), corrected for multiple comparisons at the cluster-level (*P_FWE_* < 0.05, non-parametric permutation test (first cluster (black): *P_FWE_* = .035, two-tailed; second cluster (grey): *P_FWE_* = .027, one-tailed). Horizontal lines indicate chance level decoding accuracy (.33). C) Spearman correlations between average behavioral accuracy in Component 6 experience probes and per participant averaged classification accuracy across all timepoints, post-image onset, where there is a significant difference between Condition 4 and Condition 6 in classification accuracy (see panel B). Each datapoint represents an individual participant. Condition 6 (*rho* = .48, *p* = .016) showed a numerically stronger positive correlation than Condition 4 (*rho* = .06, *p* = .787, correlation difference: *Z* = 1.53*, p =* .127, Fisher *r* to *z*). D) Spearman correlations between average behavioral accuracy in Component 6 inference probes and per participant averaged classification accuracy across all timepoints post-image onset where there is a significant condition difference between Condition 4 and Condition 6 on classification accuracy (see panel B). Each datapoint represents an individual participant. There was no evidence for a correlation difference between Condition 6 (*rho* = -.01, *p* = .969) and Condition 4 (*rho* = .14, *p* = .515, correlation difference: *Z* = .48*, p =* .630, Fisher *r* to *z*).

In line with our hypothesis, following onset of a post-transition compound stimulus presentation, classification accuracy differences were evident for participants who had experienced different graphs during prior learning (6-path graph and 6-bridge graph), spanning two distinct temporal clusters between 135 and 350 ms post stimulus-onset (Figure 4B, first cluster (black bar): cluster-level corrected *P*_FWE_= .035, two-tailed; second cluster (grey bar): *P*_FWE_ = .027, one-tailed, non-parametric permutation tests). Thus, enhanced neural decodability was evident for stimuli that shared transition types in the Component 6, but this effect was absent in participants who had experienced the 6-path graph (Component 4). This is consistent with emergence of neural similarity for stimuli that adhere to the same structural dynamics across experiential contexts, supporting a more fine-grained prediction of a representational reuse account.

To further test the robustness of the above effect, and to assess whether abstract dynamical roles are reflected in neural representational geometry during task performance, we conducted a complementary cross-phase representational similarity analysis. This analysis addresses directly whether neural representations of central nodes are stable across their distinct serial positions within a sequence, an expected property if representations reflect abstract structural identity rather than time-in-sequence. Specifically, for each participant we estimated neural patterns for each central node separately at its first and second occurrence (phase) within the sequence traversal, using occurrence-specific GLMs, and computed cross-phase similarity between prior and transfer learning phases. This approach controls for a potential time-in-sequence confound inherent in role-based comparisons, since now the same functional node is compared across different serial positions (rather than comparing different stimuli with a common sequence position, by virtue of sharing role). Consistent with the decoding results, we observed increased cross-phase similarity in participants with prior experience of Component 6 relative to those with prior experience of Component 4 (Supplementary Figure 4). Specifically, we observed a significant group difference at a late stage of the trial (1.125–1.25 sec post-stimulus onset, P_FWE_ = 0.043, two-sample non-parametric permutation test, one-tailed), with additional differences at earlier timepoints that did not survive correction for multiple comparisons (0.125 sec: t = 2.18, p = .034; 0.79 sec: t = 2.84, p = .007). These effects indicate increased neural similarity for central node representations across phases for participants who experienced these roles consistently during prior learning. We consider these results complementary to the main analyses and interpret them with caution given that the group difference was only corrected for multiple comparisons at the late cluster. Importantly, the temporal profile of the cross-phase similarity effect does not perfectly overlap with the time windows identified in the original dynamical role decoding analysis, which limits the strength of the convergence argument. We therefore cannot rule out completely a possibility that decoding effects reported in Figure 4 partly reflect temporal position within the sequence rather than abstract structural role coding per se. We additionally note that this analysis is based on a smaller and less variable set of comparisons than either the original decoding analysis or the role-based RSA, since the bridge graph contains only two central nodes whose repeated occurrences can be directly compared in this way, limiting its statistical power in isolation. Nonetheless, the observation of a group-specific effect in the cross-phase analysis which directly controls for serial position provides support for the representational reuse account, and the late corrected cluster suggests that role-consistent representations do emerge across learning phases, albeit potentially at a later stage of processing than the decoding results alone would suggest.

While our data does not provide for an unequivocal interpretation of these dynamical roles, one possibility is that individuals differentiate transitions along Component 6 based on how they relate to the unique central “bridge” formed between the 2 central nodes (e.g., the transitions between nodes 7 and 10 in Fig 3B). For example, transitions can be characterized as movement outside bridge nodes (1), arriving to/moving away from bridge nodes (2) and crossing the bridge (3). This representation of dynamical roles might confer benefits to task performance – a possibility we tested next.

### Dynamical role reuse and behavioral performance

Having found neural evidence consistent with dynamical role representations during transfer, we next asked whether the strength of these representations relates to participants’ ability to successfully reuse subprocess knowledge within new contexts. Such a relationship, if present, would provide support for the behavioral relevance of these representations. This analysis investigates whether dynamical role classification accuracies (Fig. 4B) are related to behavioral performance, and is intended as exploratory rather than providing primary evidence. Averaging classification accuracy over time points that showed significant condition differences (Fig. 4B), for experience probes we found a positive correlation between average classification accuracy and performance for Condition 6 (Condition 6: ρ = .48, p = .016, Spearman correlation; Condition 4: ρ = .06, p = .787; correlation difference: Z = 1.53, p = .127, Fisher r to z; Fig. 4C). No such relationship was evident for inference probes (Condition 6: ρ = –.01, p = .969, Spearman correlation; Condition 4: ρ = .14, p = .515; correlation difference: Z = .48, p = .630, Fisher r to z; Fig. 4D). These findings are consistent with, though not conclusive evidence for, dynamical role representations conveying behavioral benefits. Speculatively, such knowledge representation might enable individuals to constrain their predictions of successive states. For example, upon transitioning to a bridge node one can predict that the successor state will involve “the other side of the bridge”. While we acknowledge that the correlation results appear inconsistent across experience and inference probes, it is important to consider that our MEG classifiers were trained and tested on experienced transitions alone, such that enhanced classification accuracy is more reflective of a neural representation of experienced transitions. Under this assumption, the MEG classifier would be expected to align more closely with behavioral accuracy for experience probes (testing actually experienced transitions) than inference probes (testing never experienced but only implied transitions on the graph). See Discussion for further elaboration of this point, including why this asymmetry is expected under a model of gradual, imperfect structural learning.

The observed correlation is also consistent with a general learning-ability account, whereby higher-ability Condition 6 participants better discover a Component 6 structure (that happens to be shared) across sessions as well as show higher behavioral accuracy overall. For example, higher-ability participants may rediscover this structure during transfer, without necessarily benefitting from having been exposed to it during primary learning. This ability-based account predicts that decoding strength should also relate to Component 4 accuracy, and that Component 4 and Component 6 accuracy should correlate within each prior learning group. Neither prediction was supported (Supplementary Figure 5): In Condition 6, Component 6 decoding strength and Component 4 accuracy correlation were not significant (ρ = .38, *p* = .071), and did not differ significantly from the decoding strength and Component 6 accuracy correlation (Steiger’s test, *p* = .656); accuracy correlations between Component 4 and Component 6 were significant in only one prior learning group for each probe type, with no significant between-group differences (all *p* > .29).

Given this inconsistent pattern, we consider the general-ability account unsupported, though our low per-condition sample size limits power to rule it out. We therefore treat the brain-behavior correlation as supportive rather than primary evidence for a link between role representations and reuse-related performance on experience probes, pending independent replication.

Finally, we tested whether sequential memory reactivation (replay) during rest periods could account for subprocess decomposition, using temporally delayed linear modelling (TDLM) of both compound and factorized sequenceness. We did not detect reliable above-permutation-threshold sequenceness effects during any rest period, for either compound stimuli or individual features (see Supplementary Results for full details and discussion of potential limiting factors).

## Discussion

The philosophical adage of Heraclitus, “you can never step into the same river twice”, reflects much everyday experience of the world. Yet even the most complex experiences tend to share structural similarity with the past, where this often entails a recurrence of experiential subprocesses. These recurrent world properties imply that exploiting regularities derived from past experience should enable agents to effortlessly master the relentless demands of complex and ever-changing environments (Gentner, 1983). More specifically, it has been proposed that agents reuse abstract components (“building blocks”) of past experience as a “scaffolding” to facilitate integration of new information (Al Roumi et al., 2021; Luettgau et al., 2024; Pesnot Lerousseau & Summerfield, 2024). Indeed, growing evidence consistent with a compositional (re-)use of acquired knowledge provides a plausible cognitive mechanism to explain how humans transfer knowledge from prior experiences (Barron et al., 2013; Lake et al., 2015, 2017; Luettgau et al., 2024).

Here, we replicate previous behavioral findings for parsing of subprocesses that generate the dynamics of experience (Luettgau et al., 2024), and now provide neural evidence consistent with representational reuse. The neural realization of such a representational reuse is the subject of computational accounts of hippocampal and entorhinal cortex function, where grid cell activity is proposed to provide a basis set representation of abstract structure inferred from sensory experiences (Behrens et al., 2018; Mark et al., 2020; Whittington et al., 2020, 2022). However, the evolution of such structure mappings have remained elusive.

We hypothesized that success in our task is partly achieved by abstracting relational units of prior experience (’building blocks’) that bias and facilitate learning in new contexts, complementing rather than supplanting direct experience. In principle, this could be enabled by mapping new incoming sensory information onto pre-existing structural templates. We conducted three empirical tests of this representational reuse hypothesis. Addressing our first research question, whether stimuli sharing the same abstract subprocess come to be represented more similarly following learning, and whether this is specific to the subprocess experienced during prior learning, we show task-induced, condition-specific changes in neural similarity of unrelated stimuli, presented across different task phases and embedded in novel complex dynamics, conditional on these adhering to identical subprocesses and mapping onto specific structural roles. Specifically, we observed a condition-specific, learning-induced, increase in neural similarity between stimuli mapping to the same abstract structural subprocesses around 300 – 680 ms post-stimulus onset, a finding consistent with structured temporal relationships shaping neural representations across contexts (Baram et al., 2021), and with a modest degree of task structure reuse and generalization. Here, we extend these findings by detailing the neural representations associated with reuse of structural knowledge. The representational change signal observed during stimulus processing (peaking around 500 ms post-stimulus onset), echoes previous findings showing emergence of an abstract structural representation purported to account for aberrant inference and delusions in psychopathology (Nour et al., 2021). While this similarity change is consistent with neural state space alignment (Sheahan et al., 2021), it is also predicted by a more parsimonious categorical grouping/labeling of different stimuli belonging to the same abstract structural subprocess.

Addressing our second research question, whether neural representations reflect the fine-grained dynamical roles stimuli occupy within a subprocess, we found that stimuli sharing the same dynamical role across different learned structures come to exhibit neural similarity, consistent with a more specific prediction of the representational reuse account that goes beyond generic subprocess grouping. Decoding strength for these role representations showed a positive correlation with experience probe behavioral performance for the structurally relevant graph. While the direction of this brain-behavior correlation is consistent with a representational reuse account, the strength of this relationship relative to an alternative graph (4-cycle) did not reach statistical significance, and we caution this result needs independent replication to ascertain its robustness and specificity. We also acknowledge that such a brain-behavioral correlation was not present for inference probes. However, we caution that the MEG classifiers were trained and tested on experienced transitions alone such that classification accuracy reflects a neural representation of experienced transitions. Considering this, we reason that the MEG classification signal would more closely align with the behavioral accuracy in the experience probes (testing actually experienced transitions), instead of inference probes (testing never experienced but only implied transitions on the graph). We caution that the increased neural similarity we report for stimuli sharing a subprocess is also compatible, in principle, with a simpler categorical grouping account, whereby stimuli belonging to the same subprocess become more similar without encoding their specific relational role. Our role-based RSA and dynamical-role decoding analyses, which show representational change specific to stimuli occupying matched graph-theoretic roles rather than a uniform similarity across all same-subprocess stimuli, speak against a purely categorical account. We note here that this evidence is correlational and based on a single experimental paradigm.

Addressing our third research question, whether the strength of identified neural representations relate to behavioral reuse, we found a significant positive relationship between decoding strength and experience probe performance for the structurally relevant (6-bridge) graph but not for the irrelevant (4-cycle) graph. Although the relationship was numerically stronger for the relevant graph, the difference between the two relationships did not reach statistical significance. We therefore cannot establish statistically that this relationship is selective to the structurally relevant graph, and present the finding as suggestive rather than conclusive. Likewise, we did not find a corresponding relationship for inference probes. Thus, a behavioral accuracy group difference in predicting successive states for the 6-bridge inference probe performance was of negligible magnitude and did not align with the robust effect observed for experience probes (Fig. 2D), where the latter replicates and extends a previous finding using a 6-state cycle graph and predicted by computational simulations using successor feature models (Luettgau et al., 2024). We speculate that the disparity between probe types is likely due to experience probes and inference probes engaging dissociable cognitive processes – the former relying on predictive representations acquired from experience, the latter relying on an ability to perform mental operations on learned representations of independent feature transitions. This may increase the likelihood of errors due to complexities of the inferred state spaces. However, the observed differences could also relate to previously reported effects of explicit knowledge on performance levels (Luettgau et al., 2024) or be driven by capacity limitations in working memory taxing larger subprocesses more strongly (Collins et al., 2017; Collins & Frank, 2012). It is also likely that structural knowledge is not acquired uniformly across compounds.

Because abstract roles are learned gradually and with uncertainty, role inference is inherently context-dependent: the role of a given stimulus is not inferred in isolation, but in relation to the compound within which it appears. When one element of a compound is already known with high confidence, the role of the second element can be inferred more reliably; when both elements are themselves uncertain, inference becomes less precise. Learning precision may therefore vary across stimulus pairings depending on their frequency, co-occurrence statistics, and position within a sequence. On this basis we would expect corresponding variability in participants’ ability to infer dynamical roles, and hence to predict outcomes, across different compound types. This implies that an MEG signal, which indexes role-based representational structure specifically for experienced compounds, most directly reflects knowledge precision for those items. A correlation with inference-probe accuracy requires an additional inferential step, from experienced to novel compounds, introducing noise proportional to the uncertainty in the learned representations. In principle, better structural knowledge about experienced compounds should support better generalization to novel ones, but we might be insufficiently powered to detect this weaker relationship. We therefore interpret an absence of a significant correlation for inference probes not as inconsistent with representational reuse, and suggest that a null result here should be interpreted with caution rather than as evidence against it. More broadly, an absence of a strong correlation between aggregate neural decoding measures and aggregate inference-probe performance should not itself be taken as evidence against abstract learning. Inference performance is likely supported by multiple sources of information, including partial structural knowledge and heuristic strategies, whereas neural decoding captures the stability of representational geometry at the population level. The localizer RSA results reported above demonstrate that abstract role representations are present in the neural signal, even though their task-dependent expression may not always be directly mirrored in inference-probe behavior.

At a minimum our findings indicate clear learning, reuse and generalization abilities based on a factorization of complex experiences into their underlying structural elements. This entails a parsing into distinct subprocesses, derived from past experience, that are consistent with a representation of the dynamical roles these features play within distinct subprocesses. These representations, in turn, may support prediction of successive features leading to more accurate performance. Our neural findings suggest that building and maintenance of representations of subprocesses may allow factoring out the idiosyncrasies of the complex dynamics of experience. While the principal aim of the present study was to characterize neural representations consistent with a representational reuse account of structural knowledge, it is important to consider that MEG recordings reflect an aggregate neural signal. Consequently, our analyses do not support inferences about regional brain sources of the observed signal. A prior literature indicates this type of mapping relates to activity of entorhinal cortex grid cells, where these compute invariant representations of relational structure underlying experience that is abstracted away from its sensory specifics (Behrens et al., 2018; Kazanina & Poeppel, 2023; Mark et al., 2020; Whittington et al., 2020, 2022) This representational form significantly reduces a learning computational burden by lessening a requirement for inferring the structural components determining experiential dynamics, allowing an attentional focus on the idiosyncrasies of an experience (e.g., which stimulus features are currently behaviorally relevant).

The proposed representational reuse account also draws on ideas that organizing information within structured representations, including abstract neural geometries, facilitates few-shot concept learning (Sorscher et al., 2022). The neural signatures we identify are consistent with the emergence of low dimensional representations of abstract disentangled task variables, leveraged for reuse to new contexts, as previously documented in the hippocampal-entorhinal system (Baram et al., 2021; Bernardi et al., 2020; Courellis et al., 2024; Nieh et al., 2021), frontal cortex (Bernardi et al., 2020; Flesch et al., 2022, 2023; Zhou et al., 2021) or posterior parietal cortex (Flesch et al., 2022, 2023). Furthermore, evidence indicates that an encoding of abstract task structure (progress to goal), within rat medial frontal cortex, facilitates learning of complex behavioral sequences (El-Gaby et al., 2023) and that such representations emerge in artificial neural networks trained to perform multiple tasks (Johnston & Fusi, 2023; Yang et al., 2019). One implication of our findings is that representational reuse helps to align neural state spaces that encodes structure across experiential contexts. Within this account, similar state transition dynamics and dynamical roles encountered in distinct experiential contexts, results in stimuli that adhere to these being represented closer in a neural activity space (Bernardi et al., 2020; Courellis et al., 2024; Sheahan et al., 2021), putatively constraining a hypothesis space over abstract structural components that make up new experiences.

An important avenue for future research relates to the neural mechanism that supports an algorithmic implementation of the proposed representational reuse and how subprocess dynamics are disentangled. One plausible mechanism involves a separation of subprocess representations via preferential reactivation at different positions in the theta cycle, or by reactivation coupled to distinct peaks or troughs in theta oscillations (analogous to a role of theta oscillations in vicarious trial-and-error computations, Redish, 2016). Our findings hint at such a mechanism in that we show successor predictive representations are encoded neurally during transfer learning, based on decoding evidence of successor features within a current state’s neural activity pattern (Supplementary Figure 2 and Supplementary Results). Alternatively, gradual learning, and insight-dependent decomposition of an experienced compound stimulus sequence into underlying subprocesses could be enabled by human replay, extending previous accounts that propose replay has a role in compositionality (Kurth-Nelson et al., 2023; Schwartenbeck et al., 2023). We tested this directly using TDLM-based sequenceness analyses during rest periods but did not detect reliable replay effects within the constraints of our experimental design (see Supplementary Results for full details), leaving this as an open question for future work. Relatedly, there is the question of how, and under what conditions, the structure of previous experiences map to current experience. We concede that the present experiment examines reuse across two families of graph structures, rather than across a broader space of possible graphs. While we consider it unlikely that our findings are idiosyncratic to cycles, lines and bridge-like graphs, we acknowledge that further work will be needed to support abstraction beyond the limited graph structures studied here. In future experiments structural similarity and size of the reused state spaces of subprocesses in prior and transfer learning phases could be systematically varied to delineate the boundaries of mapping experiential dynamics across contexts. Future neuroimaging (fMRI) studies based on representational similarity analysis (e.g., Luettgau et al., 2022) or repetition suppression (e.g. Barron et al., 2016; Garvert et al., 2017, 2023; Luettgau et al., 2020) could also contribute to pinpointing the precise functional anatomy, including a predicted involvement of the hippocampal-entorhinal system (Baram et al., 2021; Bernardi et al., 2020; Courellis et al., 2024; Nieh et al., 2021).

A key contribution of our study is that it provides an empirical test of whether neural representations consistent with an abstract representational reuse account accompany decomposition and reuse of structural knowledge across learning contexts. Our findings are consistent with a proposal that reuse of previously abstracted relational units could bias learning and shape neural representations in novel contexts. However, we have not directly tested whether these representations causally facilitate learning or reuse, and our evidence speaks most directly to the reuse of individual components rather than to their recombination.

Our findings characterize neural representations consistent with this proposed representational reuse account, indicating that abstract structural knowledge is expressed during ongoing stimulus processing and is shaped by prior structural experience. Under this account, similar state transition dynamics encountered in highly distinct experiential contexts are represented more closely in neural activity space than dissimilar ones, allowing a sharing of components across distinct experiences. Finally, we note that failures in efficient knowledge decomposition, involving acquisition and reuse of abstract neural representations might impact a flexibility of cognition that is considered one of the hallmarks of mental health (Huys et al., 2016; Story et al., 2024).

## Methods

### Participants and procedures

Participants were recruited from the local student community at University College London (UCL) via an online recruitment platform between October 2022 and May 2023. Only participants reporting the absence of past or present mental health conditions or neurological conditions were included.

A total of 61 participants were recruited with 51 included in the final behavioral data analysis sample (*N* = 1 discontinued the experiment due to a headache, *N* = 1 excluded due to a technical error, *N* = 1 excluded due to indicating a diagnosis of aphantasia in post-experimental debriefing, *N* = 3 due to repeatedly falling asleep and closing their eyes during the experimental task, *N* = 4 were excluded due to low motivation and indicating that they gave up learning the sequence during the experiment). One additional participant was excluded from MEG data analyses due to a metal artifact, leaving 50 healthy volunteers for the final MEG sample (median age: 23 years, *SD* = 3.69, range = 18 – 36, 9 males), *N* = 25 subjects per condition. We did not perform a formal power calculation to determine the sample size but instead opted to base the sample size on that used in similar MEG studies previously conducted in our lab. Participants received £10.00 per hour as compensation. The study was approved by the University College London Research Ethics Committee (reference number: 3090/004) and conducted in accordance with the Declaration of Helsinki.

After providing informed written consent subjects were prepared for the MEG scan. Participants received written instructions regarding the pre-task localizer session on the presentation screen in the MEG scanner room. Following these instructions, subjects were asked to explain the pre-task localizer. Subsequently, participants received instructions regarding the sequence learning task and then were randomly assigned to one of the two experimental conditions (see Experimental design and behavioral task). Upon completion of the sequence learning task, participants performed a post-learning localizer task. At the end of the experiment participants filled out a sociodemographic questionnaire and a post-experimental questionnaire that assessed strategies used as well as their general understanding of the task, followed by a debriefing regarding the aims and motivation behind the study.

### Experimental design and behavioral task

#### PRE localizer

Stimuli, representing states in the graphs, comprised depictions of everyday objects, landscapes, and animals/humans, arranged to create meaningful holistic scenes (e.g., a boy and a ball, or a cat in a car). Images were selected from Pixabay (https://pixabay.com/), where two images were manually assembled to form one compound image using Inkscape (https://inkscape.org/). We used two non-overlapping and unrelated sets of ten images to compose the compound images. The assignment of sets and the positions of the two images used to create the compound images on the underlying graph structures producing the observed sequences were counterbalanced across subjects and conditions. This mitigated a potential confound in learning, as well as choice biases, due to idiosyncrasies inherent in the compound images. The PRE localizer task comprised 5 blocks, involving presentation of 960 or 1020 stimuli (depending on whether participants were allocated to the 4-cycle prior or the 6-bridge prior condition) in the center of the MEG presentation screen (stimulus presentation duration: 650 ms), with an inter-stimulus interval of 583 (±116) ms, drawn from a truncated, discretized gamma distribution (shape = 1.5, scale = 0.5), marked by a blank screen. The 20 distinct features presented during prior and transfer learning phase (10 features per task phase, respectively) with the 12 or 14 compound images (4-cycle prior or the 6-bridge prior condition, respectively) shown during the prior learning phase being presented 30 times each. Each compound image was presented as foreground to a unique background image (textures) that was randomly assigned to each compound image per participant. This design feature was intended to amplify the dissimilarity between compounds that might share individual features. Participants were instructed to monitor the presented stimuli carefully. Attention to the presented stimuli was assessed based upon 10% pseudo-randomly selected stimulus presentations being used to probe subjects regarding the last presented image. Here, a probe question, “Previous image?” was presented alongside two English words, on the left- and right-hand side to the center of the screen, describing the correct last stimulus or an alternative incorrect stimulus (order randomized across stimulus presentations, duration: 4000 ms). The words describing compound images were selected such that the holistic nature of the scene (instead of the distinct features) was emphasized. Each stimulus (features and compounds) was probed equally often. Participants responded by selecting the left or right response button and if they did not respond within 4000 ms a time out warning message was presented for 1500 ms. Participants were not provided with feedback about the accuracy of their responses but received summary feedback about their overall accuracy during the last block.

#### POST localizer

The POST localizer task was identical to the PRE localizer task. However, only the 20 features were presented, but not the compound images shown during the prior learning phase. Thus, during the POST localizer participants were presented with 600 stimuli.

#### Sequence learning task

In a sequence learning paradigm (Luettgau et al., 2024) subjects were presented with sequences of compound images. Compound images were constructed by overlaying two independently varying visual features drawn from separate feature sets. Each feature set corresponded to a single graph factor, such that the state of each graph factor independently determines one feature of the compound image. For example, a compound image in state (A, 3) would display feature A from the cyclic factor and feature 3 from the path factor, superimposed or juxtaposed into a single image.

The set of all compound images and their transitions was defined by the Cartesian product of two graph factors: transitions between compound images were valid if, and only if, one factor alone changed state according to its own transition rules while the other factor remained fixed.

Observed sequences of image compositions composed from 10 individual features. Each sequence consisted of 8 compound images with 12 compound images in total across sequences. In a between-subjects design, during prior learning one group of participants (condition 1, 4-state cycle prior) observed sequences of image compositions drawn from the product graph of a cyclic graph factor with 4 states (e.g. A->B->C->D->A …) and a path-graph factor with 6 states (e.g. 1<->2<->3<->4<->5<->6, 4 x 6 factorization). For instance, in condition 1, a valid sequence might proceed: (A,1) → (A,2) → (A,3) → (B,3) → (C,3) → (C,4) → (D,4) → (D,5), where each step reflects a transition in either the cyclic (letter) or path (number) factor, but never both simultaneously.

The other group (condition 2, 6-state bridge prior) saw compound image sequences (14 compound images total) produced by a bridge graph factor with 6 states (e.g., 7->8->9->7->10->11->12->10->7, …, 4 x 6 factorization) and a path-graph factor with 4 states (e.g., E<->F<->G<->H). Participants observed 54 image sequences during this ‘prior learning’ phase. Each compound image was presented in front of a unique background image (textures) that was randomly assigned to each compound image per participant.

Subsequently, during a transfer learning phase, we tested subjects on entirely new image compositions (16 transitions between compound images from two disjoint subgraphs of the compound graph (8 unique compound images per subgraph), composed from 10 novel individual features, Fig. 1C) where these were generated from the product graph of a cyclic graph (4 states, e.g. A’->B’->C’->D’->A’…) and a bridge graph (6 states, e.g., 7’->8’->9’->7’->10’->11’->12’->10’->7’, …, 4 x 6 factorization). The presentation of these transitions alternated between two presentations of subgraph sequence 1 (Fig. 1F, left), and two presentations of subgraph sequence 2 (Fig. 1F, right). To avoid learning “impossible” transitions between the terminal element of one sequence to the first element of the other we explicitly instructed participants regarding the start of a new sequence presentation. Participants observed 48 image sequences during this transfer learning. Note that the 6-bridge graphs used in prior and transfer learning did not feature a truly bi-directional edge between the central nodes (7//7’ and 10/10’), but rather a “refractory” edge, indicating that it was not possible to transition back and forth within one step between central nodes of the graph.

The nature of the factorization allowed us to present only a subset of 12 or 14 compound stimuli, respectively. Because sequences were drawn from a subset of the product graph, only a portion of all 24 possible compound images (and their transitions) were encountered directly. Transitions between compound images could thus be: (i) directly experienced in a sequence, (ii) never presented in a sequence but implied by the factorization structure, or (iii) transitions between compound images from different subgraphs, which were never valid. The held-out set of transitions described in (ii) and (iii) between compound stimuli (partially observed during sequence presentations, partially never observed during sequence presentations) provided a context for assessing knowledge of graph factorization.

Each sequence comprised 8 compound image presentations in the center of the screen (stimulus presentation duration: 1000 ms), interleaved by an inter-stimulus interval of 750 ms, marked by a blank screen.

Following each presented sequence, two probe questions asked participants to predict upcoming states. This assessed knowledge of either of transitions between the experienced compound stimuli in the sequence (experience probes), or their ability to infer and predict never experienced transitions between compound images (generalization), where the latter are implied by the 4 x 6 graph factorization (inference probes). We reasoned that participants could only attain above chance predictions about never experienced transitions if they had encoded a factorized representation of the true generative state space underlying the observed image compositions.

In each probe question, participants first saw a randomly selected compound image from the experienced sequence, or an entirely new compound image, with a probe question “Imagine you see this image. What would be the next image?” (duration: 3000 ms). Probe questions testing knowledge of never experienced transitions between compound images did not feature unique background images. Following the probed image, two compound images – the correct next image and a lure image – were presented as choice options on the left- and right-hand side next to the screen center. Crucially, on each probe, the lure image was matched with the correct option on one of the two features (e.g., A1 -> B2 or D2), a design feature that allowed us to test knowledge of both graph factors underlying the observed sequence separately, as well as assess knowledge about never experienced transitions between compounds. Each unique compound image in the sequence (and each never experienced transition) was used exclusively to assess knowledge of one of the two graph factors, but not both. The latter was intended to control for a possibility that participants might guess and remember the correct answer for each probe question by observing the same correct option repeatedly presented alongside different, changing lure images – allowing for the inference that the consistent option must be the correct one – even without knowing anything about the compound sequence, or correctly inferring transitions between novel compounds. Each transition within each graph factor was probed equally often in both prior and transfer learning phases.

During each probe question, participants had 4500 ms to select the left or right option by using the left or right button on two button boxes held in both hands. If they failed to respond within this time window, a timeout warning was presented for 1500 ms and the probe question was coded as a missing event. Participants did not receive feedback about whether they answered correctly or not on any given probe question. Between the experience and inference probe questions, there was a blank screen for 5000 ms (inter-probe interval). After the inference probe, the sequences proceeded.

The task comprised 8 blocks, with short breaks after every 14^th^ or 12^th^ (transfer learning phase) presented sequence and a longer break (5 min) between prior and transfer learning. During these breaks, participants received feedback about their average performance in the experience probe questions up until this trial. As well as assessing whether participants entertained a factorized representation of the graph factors generating the sequence of compound observations, our transfer learning task design allowed us to test a prediction that participants form abstractions of the decomposed or factorized state space, and reuse components of the abstracted generative processes (graph factors) they had encountered in prior learning in an entirely new context, the transfer learning phase. Importantly, the experimental design also allows us to assess a second kind of knowledge use, distinct from reusing a graph factor across learning phases: participants could repurpose their knowledge about experienced transitions (as assessed by experience probes) to generalize to never experienced transitions within the graph factors (as assessed by inference probes).

### Behavioral analyses

Data were preprocessed and analyzed in MATLAB 2023b (The MathWorks, Inc., Natick, MA, USA) and RStudio (RStudioTeam, 2019) (version 4.1.0, RStudio Team, Boston, MA) using custom analysis scripts. We defined Bayesian multilevel generalized linear models (GLM), representing different processes that might have generated the observed choice data using the R package rethinking (McElreath, 2020a, 2020b). We specified binomial likelihood functions to model the (aggregated) number of correct responses distributed as the proportion of correct probes (binomial distribution parameter *p*) among the number of probes for which a response was recorded (number of probes). We employed sampling-based Bayesian inference to estimate the posterior distribution of linear model parameters comprising different intercept and slope parameters that linearly combine to form the binomial distribution parameter *p*. In the below models, µ denotes average/group level effects (“fixed effects”), α and γ denote individual or individually covarying effects (“random effects”). We coded the probed size effect as –.5 (for 4-cycle probes) and as .5 (for 6-bridge probes). A negative value of the parameter estimate indicates higher accuracy for 4-cycle probes. The probe type effect was coded similarly (experience probes as –.5; inference probes as .5). A negative value of the parameter estimate indicates higher accuracy for experience probes. The condition effect was coded as –.5 (for 4-cycle prior condition) and as .5 (for 6-bridge prior condition). A negative value of the parameter estimate indicates higher accuracy for the 4-cycle prior condition.

For each of the below models, we specified weakly informative prior and hyperprior probability distributions (as indicated in the model specifications). Models were passed to RStan (Stan Development Team, 2020) using the function “ulam” (rethinking package). We drew 4 x 4000 samples from posterior probability distributions (4 x 1000 warmup samples), using No-U-Turn samplers (NUTS; a variant of Hamiltonian Monte Carlo) in RStan and four independent Markov chains. Quality and reliability of the sampling process were evaluated with the Gelman-Rubin convergence diagnostic measure (*R̂* ≈ 1.00) and by visually inspecting Markov chain convergence using trace- and rank-plots. For all models fitted we found *R̂* = 1.00 for all parameters sampled from the posterior distribution. There were no divergent transitions between Markov chains for any of the models reported.

For model comparisons and to find evidence for the best-fitting model for the observed behavioral data, we used the Widely Applicable Information Criterion (WAIC). Parameter estimates were considered non-zero if the Bayesian credible interval (CI) around the parameter did not contain zero.

We defined a model space of four Bayesian multilevel (statistical models, GLM1 – GLM4, Eq. 1-4) generalized linear models (cf. Luettgau et al., 2022, 2024). Both experience and inference probe correct responses were analyzed in a joint model to reduce the number of tests of the same statistical hypothesis and to increase statistical power (a joint model should capture within-subject performance ability and correlations across both probe types more adequately and explicitly than two separate models for both probe types).

To test our main hypothesis, that participants factorized and abstracted structure underlying their experience during prior learning and reused an abstracted graph factor that was consistent across prior and transfer learning, we focused on transfer learning phase behavior. We expected an advantage in predicting 4-cycle graph factor transitions in the 4-cycle prior condition (vs 6-bridge prior condition), and vice versa, enhanced performance in 6-cycle graph factor transitions in the 6-bridge prior condition (vs 4-cycle prior condition). Formally, this pattern of responses should reflect in an interaction effect between condition and probed size. We therefore predicted that the best-fitting model for the observed behavioral data (as found in model comparison) would feature a non-zero interaction effect between condition and probed size, suggesting that this effect captures important unique variance in explaining participants’ choice behavior.

GLM1. Individually varying intercepts model. This model assumes that correct responses are invariant across probed sizes, probe types, and conditions.

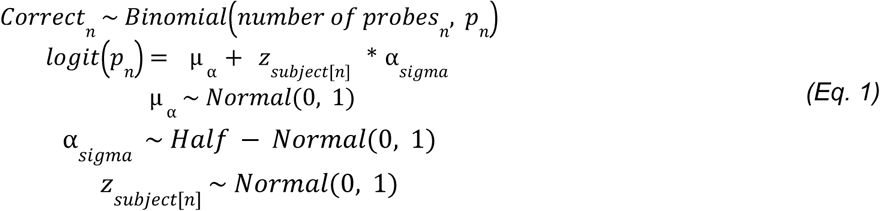

where *Correct* denotes the number of correct probes, distributed as the proportion of correct probes (binomial distribution parameter *p*) among the number of probes for which a response was recorded (number of probes), *n* denotes the parameter estimate for the n-th subject. Note that the model is reparametrized to allow sampling from a standard normal posterior distribution. α*_sigma_* denotes the standard deviation of the reparameterized normal distribution of the varying intercepts parameter α.

GLM2. Individually varying intercepts, main effect of probe type (*PT*), covarying intercept and probe type effect

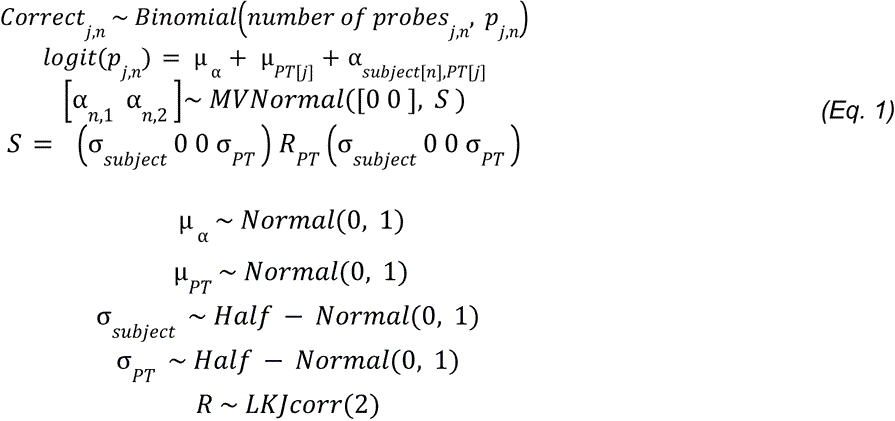

where *MVNormal* is a multivariate normal/Gaussian distribution, the correlation matrix *R_PT_* is distributed *as LKJcorr* distribution (Lewandowski-Kurowicka-Joe distribution). We used non-centered parameters for the covariance matrix *S* (Cholesky decomposition) to facilitate estimation using sampling-based inference. j denotes the parameter estimate for the j-th probe type (experience or inference probe). σ are the variance parameters for the subject specific intercept parameter and probe type parameter.

GLM3. Individually varying intercepts, main effects of probed size, probe type and condition, interaction effect of probed size x condition (*SIZE x COND*), covarying intercept and probed size, probe type effect

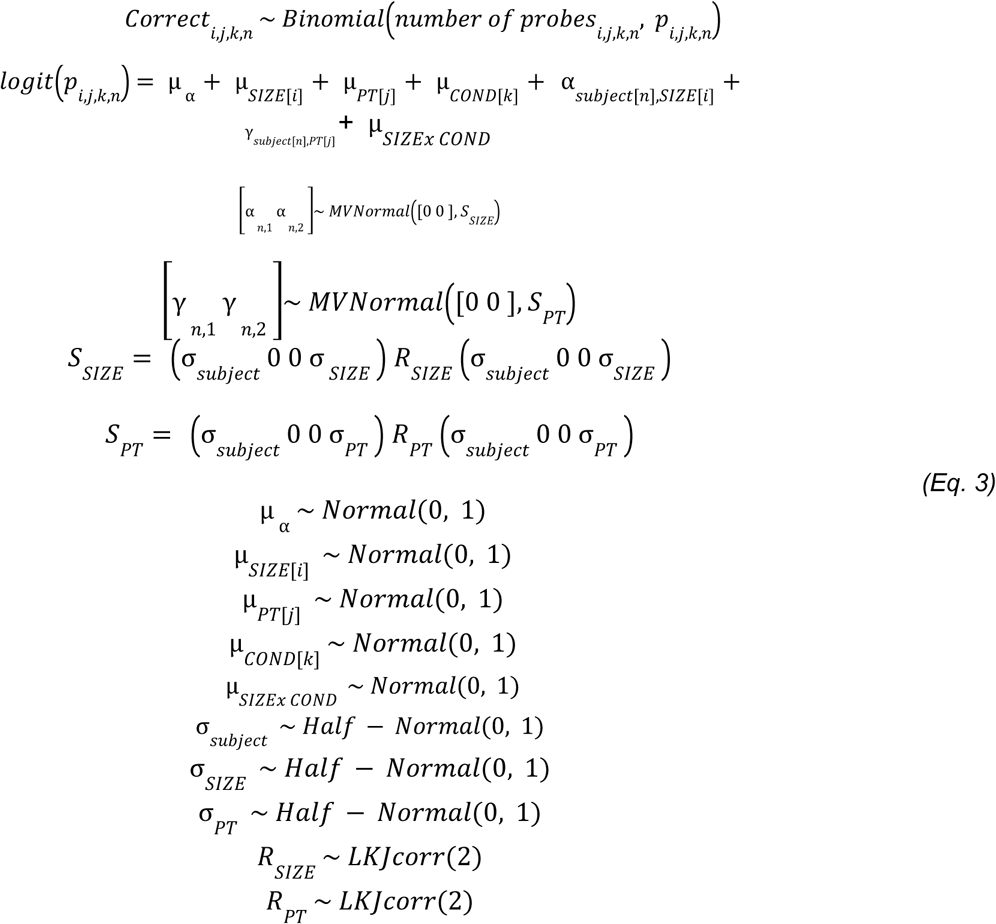

where *k* denotes the parameter estimate for the k-th condition (4-cycle prior condition or 6-bridge prior condition), i denotes the parameter estimate for the i-th probed size (4-cycle probe or 6-bridge probe) GLM4. Individually varying intercepts, main effects of probed size, probe type and condition, interaction effect of probed size x condition and probed size x probe type x condition (*SIZE x PT x COND*), covarying intercept and probed size, probe type effect

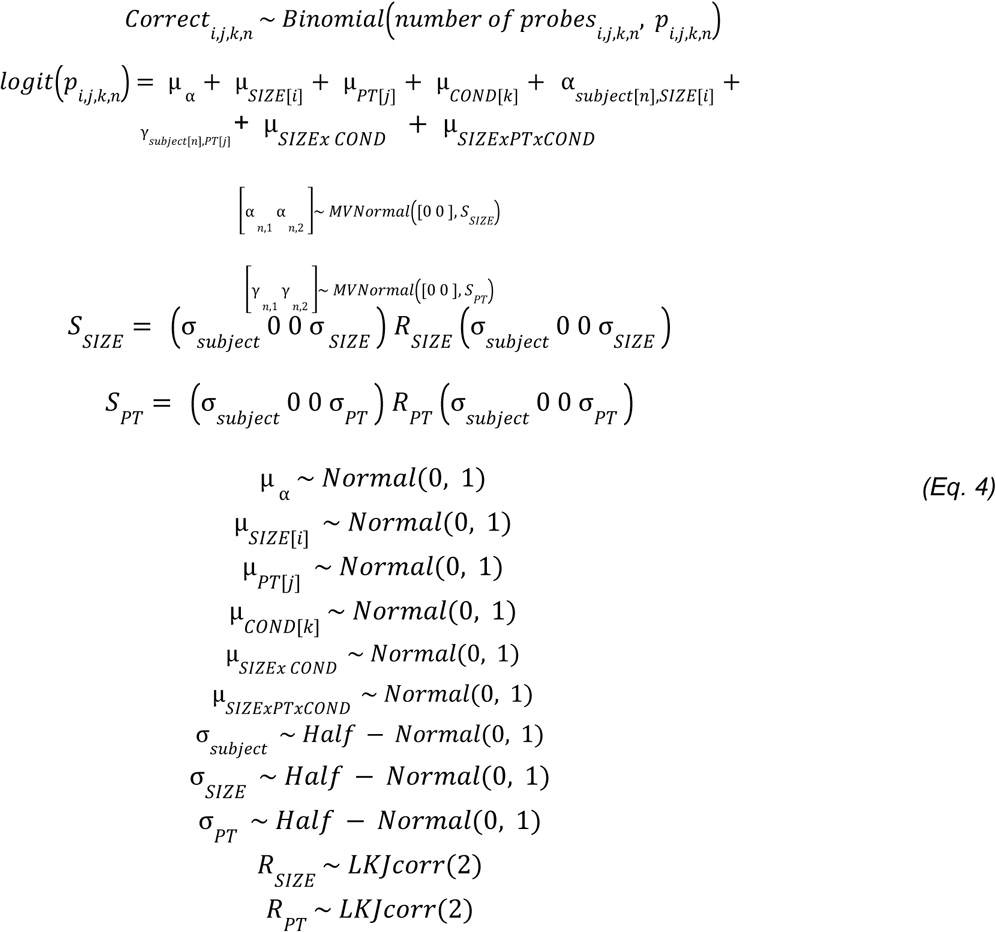

### GLM comparison

A multilevel GLM featuring all of the above effects and a three-way interaction effect of probed size x probe type x condition (GLM4) showed slightly better model fits than a multilevel GLM featuring individually varying intercepts, main effects of probed size and probe type, as well as covariation with the individually estimated intercept parameters, a main effect of condition, and an interaction effect of probed size x condition (GLM3) (WAIC values (± standard error): GLM4 = 1015.2 (±21.25), GLM3 = 1020.2 (±22.11)). All other GLMs explained the data less well than GLM4 (WAIC values (± standard error): GLM2 = 1101.4 (±33.04), GLM1 = 1151.9 (±38.60)). Crucially, there was strong model evidence that GLMs featuring varying intercepts alone (e.g., GLM1), representing the alternative hypothesis of participants entertaining a compound representation of their experience, was much worse explaining observed behavioral data.

The model comparison suggests that the effects for probed size and prior learning condition on behavioral accuracy were qualified by a three-way interaction effect between these two variables and probe type (experience vs inference) (GLM4). We thus followed up on this analysis by using post-hoc GLM3 featuring main effects for probed size, and prior learning condition and an interaction effect between these two variables separately for both probe types (excluding the probe type main effect)

### MEG acquisition and preprocessing

MEG (magnetoencephalography) was recorded at 1200 Hz using a whole-head 275-channel axial gradiometer MEG system (CTF Omega, VSM MedTech), while participants sat upright. 3 sensors were not recorded across all participants due to excessive baseline noise. Preprocessing was conducted separately for each recording block, identical to previous studies from our laboratory (e.g., Nour et al., 2021). Sensor data were high-pass filtered at 0.5 Hz to remove slow-drifts. Data were then down sampled to 100 Hz. Noisy segments and sensors were automatically detected and removed before independent component analysis (ICA). On average across all MEG runs and participants, 8.67 sensors (SD = 7.36) and 9.13 trials (SD = 11.46) were excluded (using default osl_detect_artefacts settings; channel significance level for Generalised Extreme Studentised Deviate = 0.05). We did not apply MaxFilter as our data were acquired on a CTF system, for which MaxFiltering is not designed. Sensor interpolation was not performed as bad channels are excluded prior to ICA decomposition following standard OSL pipeline practice; given that excluded sensors represented on average ∼3% of the total 273 sensors, we do not expect this to have meaningfully affected the results. ICA (FastICA, http://research.ics.aalto.fi/ica/fastica) was used to decompose the sensor data for each session into 150 temporally independent components and associated sensor topographies. Artifact components (e.g., eye blink and heartbeat interference) were automatically classified by consideration of the spatial topography, time course, kurtosis of the time course and frequency spectrum for all components (Nour et al., 2021). Artifacts were rejected by subtracting them out of the data. Before cutting the MEG data into individual stimulus presentations, we re-aligned triggers to a photodiode signal that was recorded in response to each stimulus presentation, to ensure high precision of neural time series. All MEG analyses were performed on the filtered, cleaned MEG signal from each sensor at whole-brain sensor level, in units of femtotesla. Note that for each participant different MEG sensors were excluded, based on the identified noise level. Only sensors that were not identified as noisy consistently across localizer or main task runs – depending on the respective analyses – were included. We treated noisy sensors to be missing completely at random.

### MEG data analysis

#### Feature similarity analysis

We conducted a feature similarity analysis (inspired by representational similarity analysis (RSA) (Kriegeskorte et al., 2008)) to investigate task-related changes in abstract graph factor representations from PRE to POST localizer. To this end, for each session we z-scored the pre-processed MEG data over all stimulus presentations, for each sensor and from -200 to 800 ms relative to stimulus-onset. To denoise the data and reduce dimensionality, the z-scored sensor x time x stimulus presentations data was subjected to principal components analysis (resulting in a PC [20] x time x stimulus presentations data matrix, 20 PCs per subject, explaining >= 90% of the variance). We regressed the neural data onto a session-specific design matrix, denoting the stimulus label of each stimulus presentation (dummy coded). We used the resulting [feature x 1] vector of regression weights, as an estimate of the unique PC activation related to each feature. We repeated this procedure for all PCs. We then calculated the pairwise absolute differences between features as distance metric at each time point, for each PC separately. This generated a symmetrical [20 × 20] Representational Dissimilarity Matrix (RDM) at each time point. To retain as much stimulus similarity information contained within the neural data as possible, while discarding noisy components, we used the 20 PC-related RDMs to compute a weighted average RDM for each participant at each time point (in both localizer tasks separately). The weighting of the average RDM was based on explained variance of the PCs included in the analysis. For each participant individually this encompassed the top RDM computed on the PCs that cumulatively explained >= 80% of the variance of the data. To enable an estimate of task-induced PRE-POST changes, we then computed the difference in weighted averaged RDMs (POST minus PRE). We regressed out across set confusion (e.g., prior learning 4-state graph factor and transfer learning 6-state graph factor) to control for general effects of set similarity. Finally, on the residuals of this regression, we computed Kendall’s tau to quantify the variance in feature similarity change at each time point that was uniquely explained by an abstracted representation of the graph factors (separate design matrices for the 4-state graph factor and 6-state graph factor). The feature similarity change was smoothed over time with a 25 ms Gaussian kernel prior to computing Kendall’s tau.

In addition to the PCA-based feature analysis, we performed a sensor-level RSA. At each time point, we estimated condition-specific activation patterns using a GLM applied separately to each sensor (dummy-coded stimulus identity). RDMs were then computed across stimuli using mean absolute difference (L1 norm) as the primary distance metric. For completeness, we additionally evaluated Euclidean distances (L2 norm), which yielded qualitatively similar results (Supplementary Figure 3).

To test whether localizer representations of abstract graph structure were shared across sets experienced in the main task, we performed RSA separately for PRE and POST localizers. Structural roles were derived from each participant’s learning-phase transition sequences. For 4-state graphs, nodes were assigned to two matched role bins (boundary-like vs central-like) based on first-encounter order within each phase. For 6-state graphs, roles were determined either from node degree (6-line: boundary, intermediate, central). For 6-bridge graphs we used a predefined structural template. Specifically, the first eight valid transitions in the learning phase were mapped onto a fixed sequence of structural roles (central–intermediate–boundary–central–central–intermediate–boundary–central). Each node was assigned a role based on the majority role it occupied across its occurrences in these eight steps (ties resolved in favour of central > intermediate > boundary). For the localizer data, at each time point and sensor we estimated condition-specific activations using a GLM with 20 dummy-coded regressors (10 identities × 2 sets). This yielded sensor-wise beta patterns for each condition. Cross-set representational dissimilarities were computed using an L1 distance metric (mean absolute difference across sensors), resulting in 4×4 and 6×6 cross-set RDMs at each time point. We then summarized each RDM according to structural role: within-role distances (same role across sets), between-role distances (different roles), and their difference (delta = between − within). To control for unequal pair counts, between-role comparisons were subsampled to match the number of within-role pairs (200 repetitions). This produced time-resolved structural generalization indices for 4-state and 6-state graphs, which were submitted to group-level statistical analyses as described above.

We used nonparametric tests to identify timepoints where there was evidence for a condition (4-cycle prior condition and 6-bridge prior condition) x graph factor (4-factor and 6-factor) interaction effect, based on a group-level GLM (including condition, graph factor main effects and a condition x graph factor effect), correcting for multiple comparisons over time. At each timepoint, we computed the GLM and extracted the *F*-value for the interaction effect. We then identified contiguous clusters of time points where the observed *F*-values exceeded a predefined threshold (critical *F*-value) and computed the sum of *F*-values (*F*-sum) within each cluster. To correct for multiple comparisons, we performed a cluster-based permutation procedure that involved 1000 permutations. In each permutation, we shuffled the condition labels across participants before recomputing the *F*-values for the interaction effect at each time point, extracting the largest cluster-level *F*-sum from each permutation to build an empirical null distribution. An empirical *p*-value was computed by calculating the proportion of permutations for which the empirical absolute *F*-sum was smaller than or equal to the absolute permuted *F*-sums. A cluster in the observed data was deemed significant family-wise error corrected at the cluster-level if the empirical *p*-value was *P*_FWE_ < 0.05. Similarly, we conducted permutation tests for the post-hoc tests, comparing the two conditions on the two graph factors (two-sample tests) using the same cluster-based correction approach.

#### Decoding of dynamical roles

The following analysis aimed to assess neural evidence for an abstract representation of transitions in the 6-state graph factor.

We investigated different transition types between images on the 6-state graph factor (i.e., the 6-state path-graph factor or the 6-state bridge graph factor) (Figure 4A). While the 6-state graph factor permits the categorization of transitions into three types, such differential labeling and classification is not possible for the 4-cycle graph factor, where all transitions are structurally identical with respect to the outbound and inbound node. In the 6-state graph factor, we grouped each transition between features into their three distinct types: boundary (1), intermediate (2), and central (3). Boundary transitions represent the transitions at the graph factor’s peripheries, occurring between nodes at extreme ends. Intermediate transitions represent the transitions from boundary nodes towards the center, bridging the boundary and central nodes of the graphs. Center transitions represent transitions within the two central nodes of the graph.

We analyzed neural activity in a window from 250 ms pre- to 1250 ms post-stimulus onset for each post-transition stimulus. The 1250 ms post-stimulus period encompassed the 1000 ms image display duration. Neural activity during stimulus presentations following each transition label was used to train L1-regularized logistic regressions to classify transitions during the prior learning phase (one classifier per transition type, trained in a one-vs-all fashion). Classifiers were then tested for generalization in the transfer learning phase with different stimulus sets used in both phases. No cross-validation was performed since training and testing were on independent datasets.

In the testing phase, we applied the trained classifiers to predict each transition type given the neural activity. To compute classification accuracy, at each sample time point, we selected the classifier with the highest predicted probability and compared the maximum predicted class to the correct ground truth label class.

Further, we conducted two-sample t-tests at each time point to compare classification accuracy between participants in 4-state cycle prior and 6-state bridge prior condition. To correct for multiple comparisons, similarly as described for the RSA, we performed a cluster-based permutation procedure with 1000 permutations. At each timepoint, we computed a two-sample t-test and extracted the *t*-value. We then identified contiguous clusters of time points where the observed *t*-values exceeded a predefined threshold (critical *t*-value) and computed the sum of *t*-values within each cluster. In each permutation, we shuffled the condition labels across participants before recomputing the *t*-values for the condition comparison at each time point. We then identified contiguous clusters of time points where the observed *t*-values exceeded a predefined threshold and used the largest cluster-level *t*-sum from each permutation to build an empirical null distribution. An empirical *p*-value was computed by calculating the proportion of permutations for which the empirical absolute *t*-sum was smaller than or equal to the absolute permuted *t*-sums. A cluster in the observed data was deemed significant family-wise error corrected at the cluster-level if the empirical *p*-value was *P*_FWE_ < 0.05.

Separately for each prior learning condition, we computed the classification accuracy averaged across the time points showing a significant condition difference. Subsequently, we computed the Spearman correlation coefficient for the condition-averaged classification accuracy at the timepoints of significant condition differences and the 6-bridge probe performance (separately for experience and inference probes) for each condition separately. To assess significant differences between correlation coefficients, they were transformed to z-scores and compared (Fisher’s *r* to *z* transformation).

## Code Availability

All code used to generate the results and figures in this paper will be made on GitHub upon publication.

## Data Availability

Behavioral data and MEG data used to generate the results and figures in this paper will be made available on GitHub upon publication.

## Acknowledgements

Financial acknowledgments: RJD (Wellcome Investigator Award, 098362/Z/12/Z). The Max Planck UCL Centre is supported by UCL and the Max Planck Society. The Wellcome Centre for Human Neuroimaging (WCHN) is supported by core funding from the Wellcome Trust (203147/Z/16/Z). Open access funding provided by Max Planck Society.

The authors would like to thank all volunteers who took part in the study. We are grateful for the generous support of the imaging support team at UCL WCHN, particularly Daniel Bates, Chloe Carrick, Yasmin Feuozi, and Dimitra Moraiti for thoughtful discussions during scanning times. We would also like to thank Matthew Nour for helpful discussions during the setup of the study and for sharing analysis code. We thank Oliver Vikbladh and Neil Burgess for insightful discussions of task design & results and Leo Chi U Seak for helpful comments on an earlier version manuscript. We thank the members of the Max Planck UCL Centre for Computational Psychiatry and Ageing Research and the DeepMind Neuro-Lab for discussions at various stages of the project. We additionally thank Magda Dubois for creative suggestions of words describing the compound images used in the experimental task.

## Contributions

LL, RM, ZK-N, RJD conceived and designed the study. LL and NC acquired the data. LL and NC analyzed the data. TE and SV advised on analyses. RM, ZK-N, RJD supervised analyses. LL and NC drafted the manuscript. All authors read and edited versions of the manuscript and approved the final version of the manuscript.

## Supplementary Information

## Supplementary Results

### Neural successor feature representations

Increased neural similarity for stimuli belonging to identical abstract subprocesses is consistent with reuse of relevant structure across experiential contexts. On this basis, we asked whether neural activity also carries predictive (successor feature) representations of experienced subprocess dynamics.

We focused on characterizing neural signals that reflect knowledge representation of experienced subprocess dynamics during transfer learning. If indeed neural activity encodes predictive representations, then we should be able to predict successor features from a current state’s neural activity pattern. To test this, we trained classifiers on neural activity recorded for each individually presented feature during the PRE task localizer phase (Fig. 3A, see Supplementary Methods for details). Then, as a neural metric of successor feature representation, we tested the classifiers’ generalization ability at transfer learning – generating the predicted probability for the successor features for each compound stimulus presentation, separately for Component 4 and Component 6 successor features (see Methods for details on the classification scheme). Across several time points post-stimulus onset, we found increased predicted probabilities that were above average pre-stimulus onset predicted probabilities (Supplementary Fig. 2, *N* = 49 due to convergence issues of classifiers in one participant in Condition 4) for successor features pertaining to both graph factors (cluster-level corrected *P*_FWE_= .002 and *P*_FWE_= .037, non-parametric permutation test; for Component 4 and Component 6 successor features, respectively), consistent with a neural representation of successor features for both experienced subprocesses after stimulus onset.

**Supplementary Figure 2.**
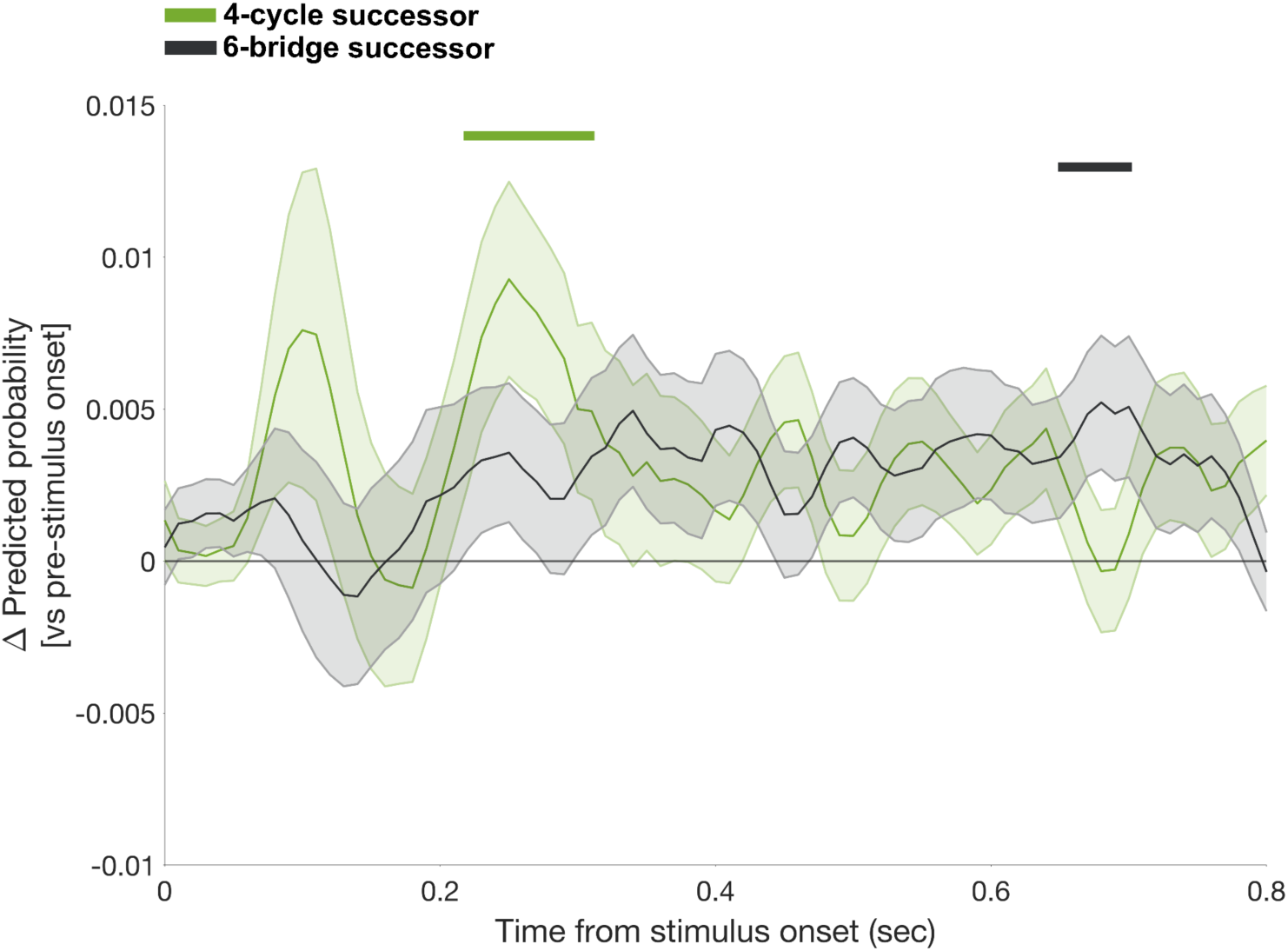
Time course of successor feature predictions. Time courses of change in predicted probability of classifiers (reactivation) relative to average pre-stimulus onset baseline during transfer learning across both conditions, related to all stimulus presentations, for the respective successor features of each compound stimulus presentation. We trained multivariate classifiers (L1-regularized logistic regressions, trained excluding the respective other feature in each compound) to distinguish between features on Component 4 and Component 6 during the PRE localizer tasks and tested the compound-by-compound successor feature predictions on both graph factors separately. Depicted are average changes in predicted probability (subtracting average pre-stimulus onset predicted probability), shaded error bars indicate standard errors of the mean. Horizontal line indicates null effect (0 predicted probability change). Solid lines on top index temporal clusters wherein the predicted probability is statistically significant above 0, corrected for multiple comparisons at the cluster-level (*P_FWE_* < 0.05, non-parametric permutation test, one-tailed).

**Supplementary Figure 3.**
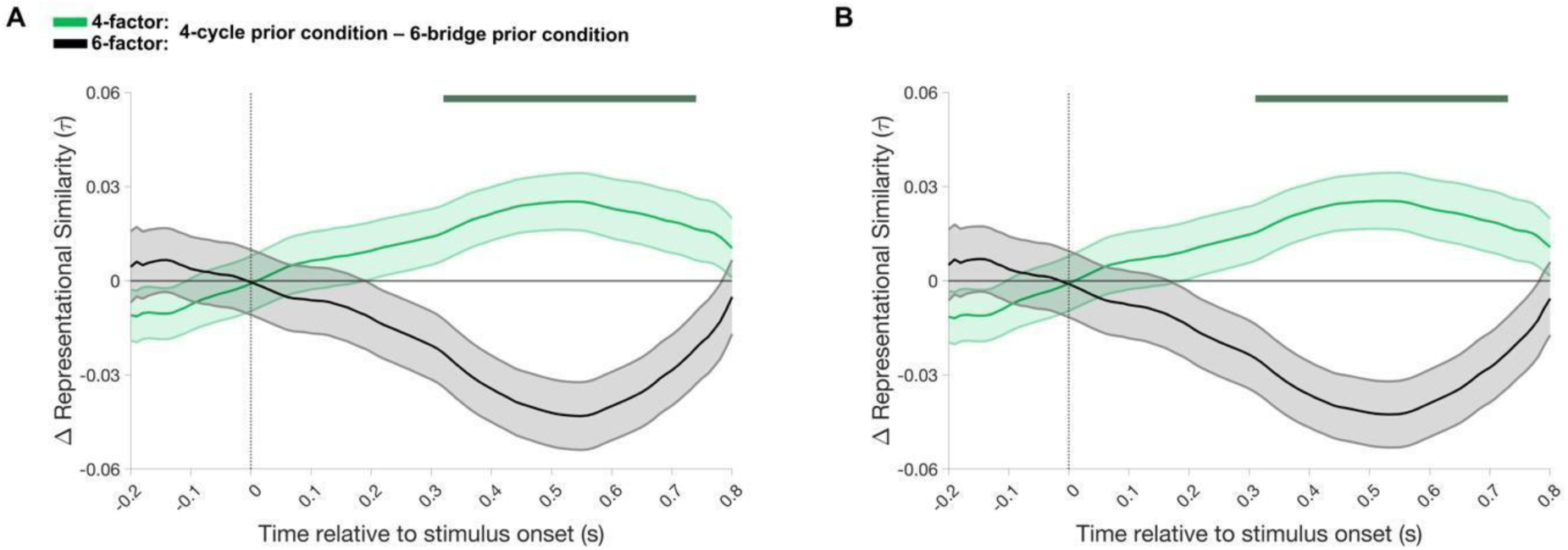
Alternative distance metrics for results shown in Figure 3D. A) Time courses of condition differences (4-cycle prior condition – 6-bridge prior condition) on Component 4 (green) and Component 6 (black) feature similarity changes from PRE to POST localizer at the sensor-level, using absolute distance (L1 norm) as similarity metric. Shaded error bars indicate standard errors of difference. Dark green line represents time points at which feature similarity changes for the condition x graph factor interaction effect were statistically significant, corrected for multiple comparisons at the cluster-level (P_FWE_ < 0.05, non-parametric permutation test). Horizontal line indicates a null effect. B) Same as A) for Euclidean distance (L2 norm) as similarity metric.

**Supplementary Figure 4.**
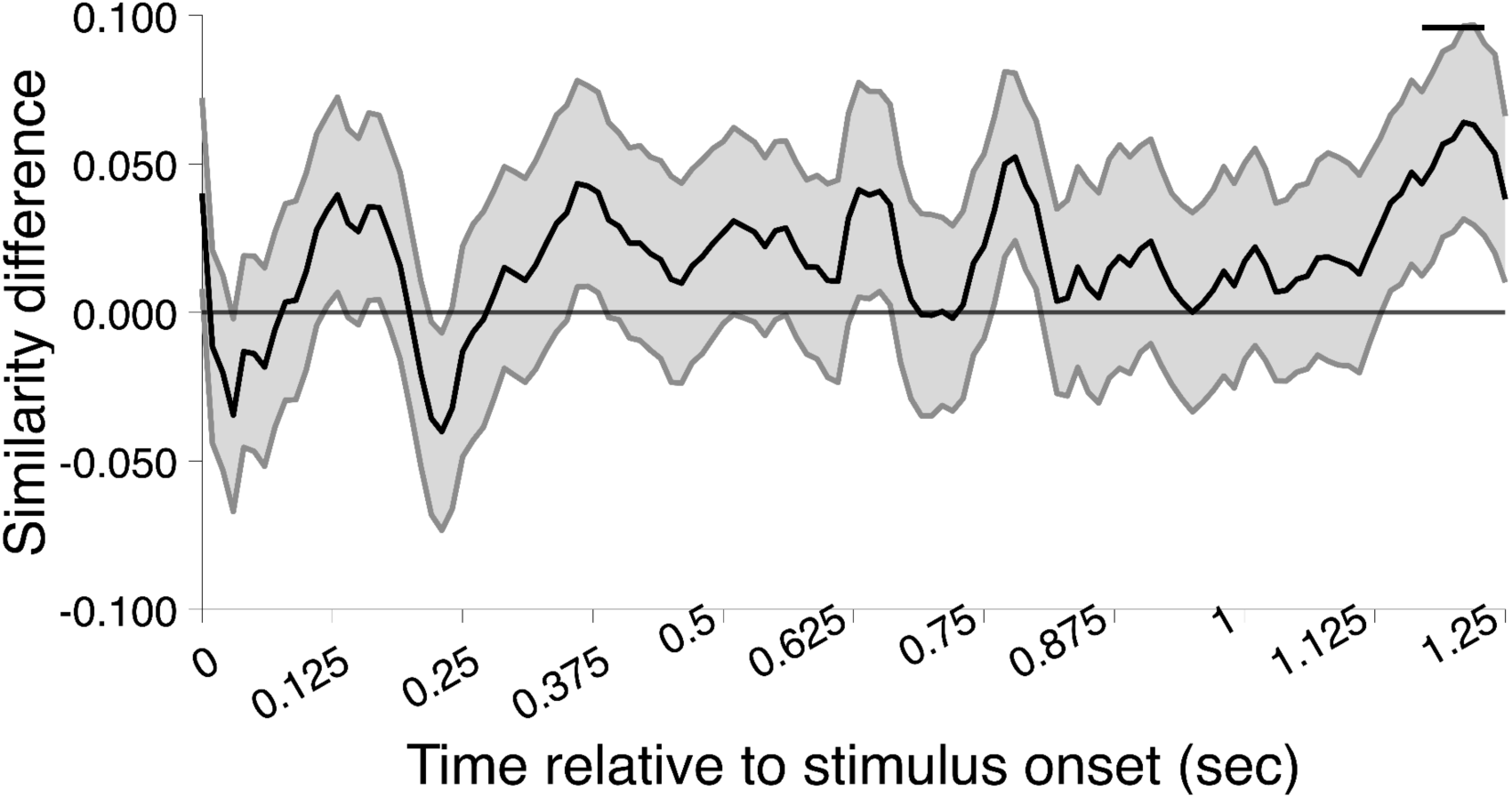
Cross-phase representational similarity of central nodes. Time courses of cross-phase representational similarity for central nodes of Component 6, computed from sensor-level data. Time courses reflect the difference in L1 distance between Condition 6 and Condition 4 (positive values indicate higher similarity for Condition 6). For each participant, central node neural patterns were estimated separately for the first and second presentation of each central node within a given sequence. Similarity was computed as the mean L1 distance across four cross-phase pairs: central node A (first presentation in prior, second in transfer), central node A (second presentation in prior, first in transfer), and the equivalent pairs for central node B. Timecourses are baseline-corrected (pre-stimulus period) and restricted to post-stimulus timepoints. The shaded error bar indicates the standard error of the difference between groups. The black bar above the time course indicates timepoints where Condition 6 differed significantly from Condition 4 (corrected for multiple comparisons at the cluster level, P_FWE_ < 0.05, two-sample non-parametric permutation test, one-tailed).

**Supplementary Figure 5.**
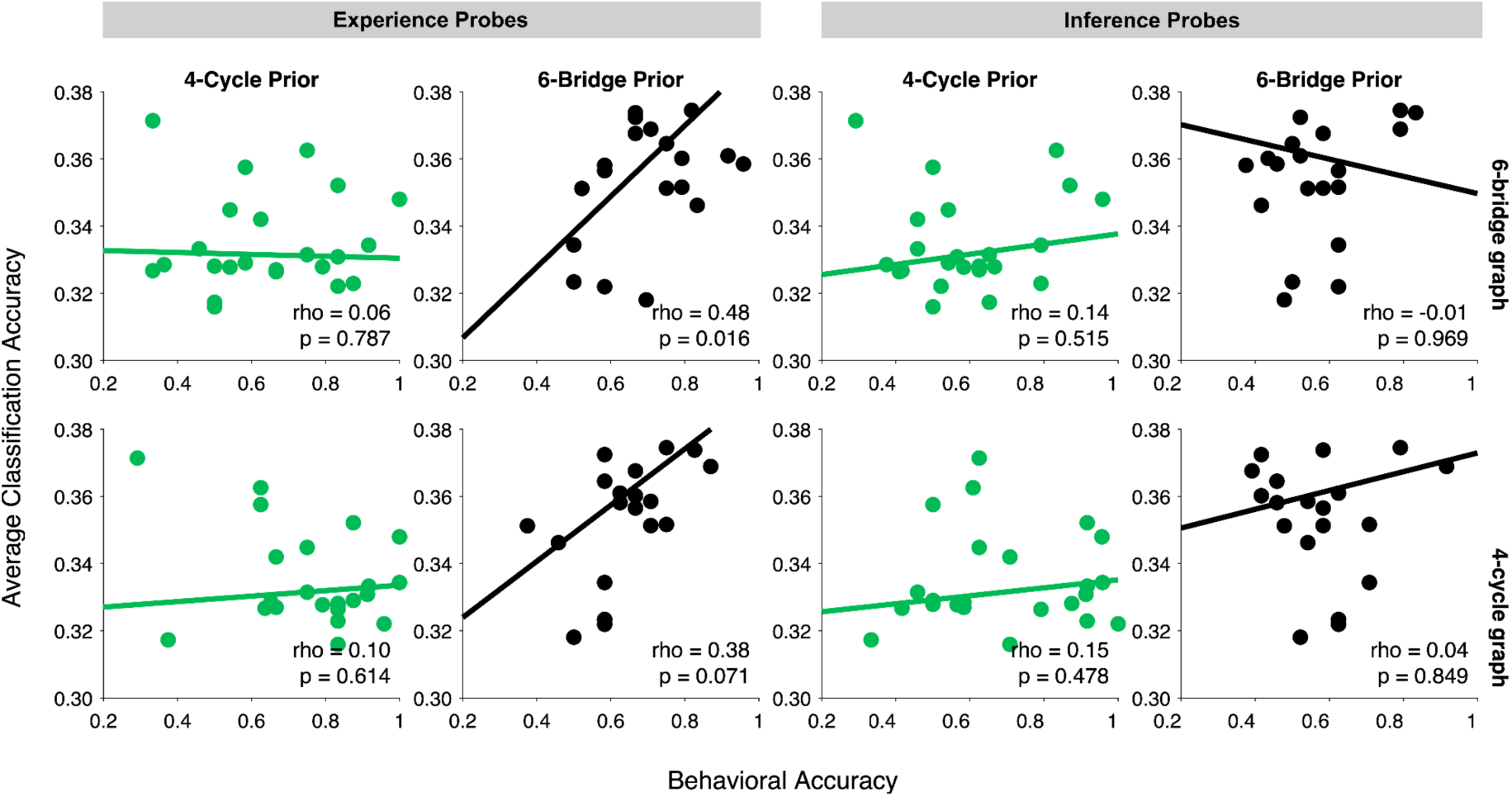
Brain–behavior correlations with Component 4 and Component 6 performance during transfer learning. Scatter plots show the relationship between average neural decoding strength, computed at timepoints exhibiting a significant group difference (Fig. 4B), and behavioral transfer-learning performance, separately for Condition 4 (green) and Condition 6 (black). Rows show performance on Component 6 (top) and Component 4 (bottom). Columns show experience probe performance (left two columns) and inference probe performance (right two columns). Each point represents one participant; solid lines show least-squares fits. Rho and p-values (bottom right of each panel) reflect Spearman correlations with permutation-based significance testing (10,000 permutations, two-tailed).

### Replay analyses

Building on work on human replay, as measured with MEG, in our and other research groups (e.g., Huang & Luo, 2024; Kern et al., 2024; Kurth-Nelson et al., 2016; Liu et al., 2019, 2021; Nour et al., 2021; Schwartenbeck et al., 2023) we tested for the presence of two qualitatively distinct types of sequential reactivation (“sequenceness”) using temporally delayed linear modeling (TDLM). Specifically, we tested for different magnitudes of sequenceness that captured transitions between neural representations of compound stimuli (“compound sequenceness”) and condition differences of sequenceness capturing transitions between neural representations of the individual features (“factorized sequenceness”). This analysis aimed at extending previous accounts of replay compositionality (Kurth-Nelson et al., 2023; Schwartenbeck et al., 2023). In these analyses we did not detect reliable, above permutation threshold sequenceness effects during a 5-minute rest period between prior and transfer learning, neither for compound stimuli nor for individual features. Such sequenceness effects were also absent during the initial 5-minute resting period before the PRE task localizer. Additionally, we did not observe any evidence for one-step transitions between neural representations of compounds or features at rest.

We also tested an additional hypothesis regarding a potential neural mechanism supporting gradual learning, and insight-dependent, decomposition of the experienced compound stimulus sequence into underlying subprocesses. Specifically, we examined for a trial-by-trial positive linear trend for factorized sequenceness and a trial-by-trial negative linear trend for compound sequenceness during the resting period between the experience and inference probe question (inter-probe intervals). Again, we did not detect significant and reliable sequenceness effects.

The absence of sequential reactivation findings in these data may relate to several factors. Firstly, we optimized hyper-parameters and classifier performance on visual localizer tasks and tested their generalization on memory reactivation during “no-task” resting periods. The L1-regularization training procedure minimizes an overlap between representations of visually presented stimuli by reducing the magnitude of regression weights for non-influential sensors. Conceivably, this might impede an ability to reliably detect memory reactivation during “no-task” resting periods, rendering quantification of sequential memory reactivation challenging. Secondly, it is possible that the reuse behavior seen in the present task may not rely on “offline” sequential reactivation of experienced task states. The behaviorally relevant information to be learned by participants encompassed one-step transitions between compound states (and features) alone, potentially rendering underpowered any sequential reactivation analyses that relies on multiple-step transitions of reactivated states.

## Supplementary Methods

**Supplementary Figure 1.**
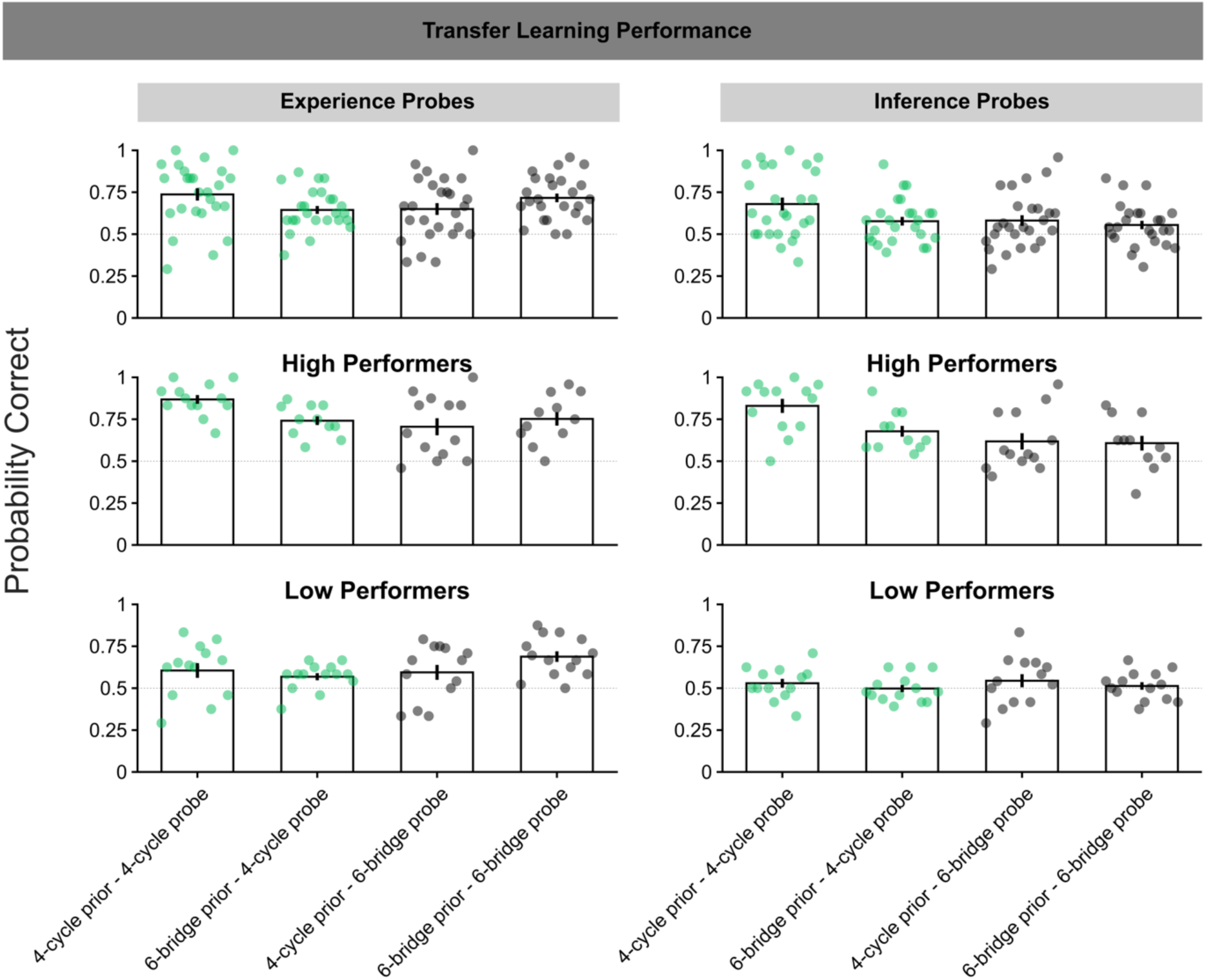
Transfer learning performance. Aggregated probabilities of correct answers during the transfer learning phase, separately for both experience (left) and inference probes (right) as a function of probed size (Component 4 (green) vs Component 6 (black)) and condition (Condition 4 vs Condition 6). Full sample (top row), high performers (>50th percentile, middle row) and low performers (<=50th percentile, bottom row). Each dot represents one participant, bars represent the arithmetic mean of the distribution, error bars depict the standard error of the mean and the dashed gray line represents chance level point estimate (probability correct = 0.5).

### Successor feature prediction

This analysis aimed at assessing evidence for neural representation of successor features in both subprocesses.

We employed L1-regularized (LASSO) logistic regression classifiers to MEG data recorded during stimulus presentations in the PRE task localizer and transfer learning phase. Individually optimized L1 penalty hyper-parameter values (λ) per participant were obtained using leave-one-run-out cross-validation (5-fold CV, training on four of the localizer blocks and testing classifier performance on the remaining block) individually on neural activity recorded at all individually presented transfer learning phase features during the PRE task localizer. Optimized hyper-parameters were used to train classifiers per participant on the transfer learning phase features presented during the PRE task localizer. Since we employed a one vs all classification scheme and every compound stimulus presented during the transfer learning phase was composed of two features, we reasoned that the classifier representing the other (irrelevant) feature in the compound stimulus would show the highest predicted probability. We sought to control for the confounding influence of the other stimulus (of no interest) contained in the compound stimulus by training and testing classifiers excluding the respective other stimulus, resulting in n * n-1 (10 * 9 = 90) classifiers. We then applied the trained classifiers to test their generalization abilities in the transfer learning phase – generating the predicted probability for each successor feature for each subprocess separately, on each compound stimulus presentation. Since all classifiers showed some level of predicted probability, we used the raw predicted probability as a neural metric of successor state representation (instead of computing the classification accuracy for the next state only). No cross-validation was performed since training and testing were performed on independent datasets.

We analyzed neural activity in a time window from 0 – 800 ms post-stimulus onset for each presented compound stimulus. This interval was chosen since the PRE task localizer presented each stimulus for 650 ms. The 800 ms post-stimulus period was shorter than the 1000 ms image display duration in transfer learning – we therefore make no claims about successor state reactivations that might have potentially occurred between 800 and 1000 ms post-stimulus onset.

To quantify the neural representation of successor features, we subtracted the averaged pre-stimulus onset (-200 – 0 ms) predicted probabilities on a given stimulus presentation from the post-stimulus predicted probabilities – providing a neural metric for feature reactivation. We conducted one-sample t-tests against 0 (null effect) predicted probability, jointly for both prior learning conditions at each time point post-stimulus onset for Component 4 and Component 6 subprocess successor feature representation. We then identified contiguous clusters of time points where the observed *t*-values exceeded a predefined threshold (critical *t*-value) and computed the sum of *t*-values within each cluster. To correct for multiple comparisons, we performed a cluster-based permutation procedure with 1000 permutations. In each permutation, we shuffled the sign of each of the successor feature activations before recomputing the t-values vs 0 at each time point. We then identified contiguous clusters of time points where the observed *t*-values exceeded a predefined threshold and used the largest cluster-level *t*-sum from each permutation to build an empirical null distribution. An empirical *p*-value was computed by counting the number of times the original cluster sum was greater than or equal to the permuted *t*-sums. A cluster in the observed data was deemed significant family-wise error corrected at the cluster-level if the empirical *p*-value was *P*_FWE_ < 0.05.

### Cross-phase representational similarity analysis

Complementary to the decoding analysis in Figure 4, we conducted a cross-phase representational similarity analysis (RSA) to test whether neural representations of central nodes were stable across their distinct serial positions within a sequence, a property expected under abstract structural role coding but not under serial position coding. We focused on central nodes of the 6-state graph, as these nodes appear twice within each sequence traversal of the bridge graph: the left central node occurs at steps 1 and 4, the right central node occurs at steps 5 and 8, allowing first and second presentations to be dissociated.

Structural roles were assigned separately for prior and transfer learning phases. For line graphs (experienced by the Condition 4 participants during prior learning), central nodes were identified from node degree in the adjacency matrix (boundary: degree 1; intermediate: neighbors of boundary; central: remaining nodes), and the two central nodes were ordered by first appearance in the trial list. For bridge graphs, central nodes were identified from the predefined 8-step traversal template.

At each time point and sensor, condition-specific activations were estimated using occurrence-specific GLMs: one GLM fitted to trials corresponding to the first occurrence of each number in the trial list, and one fitted to trials corresponding to the second occurrence. Component 6-specific beta estimates were extracted from each GLM separately, yielding four sets of neural patterns per subject: prior-phase first occurrence, prior-phase second occurrence, transfer-phase first occurrence, and transfer-phase second occurrence.

Cross-phase representational dissimilarity was then computed using an L1 distance metric (mean absolute difference across sensors) across four cross-phase pairs: Left central (prior first, transfer second), left central (prior second, transfer first), right central (prior first, transfer second), and right central (prior second, transfer first). These pairings compare the same node across different serial positions and different learning phases, directly controlling for time-in-sequence as a potential confound. The mean L1 distance across the four pairs constituted the cross-phase dissimilarity index at each time point, where lower values indicate greater neural similarity. Timecourses were baseline-corrected by subtracting the pre-stimulus mean (timepoints 1–25, corresponding to −250 to 0 ms) per participant, and analyses were restricted to post-stimulus timepoints. Group differences between Condition 6 and Condition 4 participants were assessed using a two-sample non-parametric cluster permutation test (cluster-forming threshold p = 0.10, cluster-level alpha = 0.05, 1,000 permutations, one-tailed, consistent with the directional prediction that Condition 6 participants would show greater cross-phase neural similarity for central nodes). The results are presented in Supplementary Figure 4.

## References

Al Roumi, F., Marti, S., Wang, L., Amalric, M., & Dehaene, S. (2021). Mental compression of spatial sequences in human working memory using numerical and geometrical primitives. Neuron, 109(16), 2627–2639.e4. 10.1016/j.neuron.2021.06.009

Allen, K. R., Smith, K. A., & Tenenbaum, J. B. (2020). Rapid trial-and-error learning with simulation supports flexible tool use and physical reasoning. Proceedings of the National Academy of Sciences of the United States of America, 117(47), 29302–29310. 10.1073/pnas.1912341117

Baram, A. B., Muller, T. H., Nili, H., Garvert, M. M., & Behrens, T. E. J. (2021). Entorhinal and ventromedial prefrontal cortices abstract and generalize the structure of reinforcement learning problems. Neuron, 109(4), 713–723.e7. 10.1016/j.neuron.2020.11.024

Barron, H. C., Dolan, R. J., & Behrens, T. E. J. (2013). Online evaluation of novel choices by simultaneous representation of multiple memories. Nature Neuroscience, 16(10), 1492–1498. 10.1038/nn.3515

Barron, H. C., Garvert, M. M., & Behrens, T. E. J. (2016). Repetition suppression: a means to index neural representations using BOLD? Philosophical Transactions of the Royal Society B: Biological Sciences, 371(1705), 20150355. 10.1098/rstb.2015.0355

Behrens, T. E. J., Muller, T. H., Whittington, J. C. R., Mark, S., Baram, A. B., Stachenfeld, K. L., & Kurth-Nelson, Z. (2018). What Is a Cognitive Map? Organizing Knowledge for Flexible Behavior. Neuron, 100(2), 490–509. 10.1016/j.neuron.2018.10.002

Bernardi, S., Benna, M. K., Rigotti, M., Munuera, J., Fusi, S., & Salzman, C. D. (2020). The Geometry of Abstraction in the Hippocampus and Prefrontal Cortex. Cell, 183(4), 954–967.e21. 10.1016/j.cell.2020.09.031

Collins, A. G. E., Ciullo, B., Frank, M. J., & Badre, D. (2017). Working memory load strengthens reward prediction errors. Journal of Neuroscience, 37(16), 4332–4342. 10.1523/JNEUROSCI.2700-16.2017

Collins, A. G. E., & Frank, M. J. (2012). How much of reinforcement learning is working memory, not reinforcement learning? A behavioral, computational, and neurogenetic analysis. European Journal of Neuroscience, 35(7), 1024–1035. 10.1111/j.1460-9568.2011.07980.x

Conway, C. M., & Christiansen, M. H. (2006). Statistical Learning Within and Between Modalities Pitting Abstract Against Stimulus-Specific Representations.

Courellis, H. S., Minxha, J., Cardenas, A. R., Kimmel, D. L., Reed, C. M., Valiante, T. A., Salzman, C. D., Mamelak, A. N., Fusi, S., & Rutishauser, U. (2024). Abstract representations emerge in human hippocampal neurons during inference. Nature, 632(8026), 841–849. 10.1038/s41586-024-07799-x

Dekker, R. B., Otto, F., & Summerfield, C. (2022). Curriculum learning for human compositional generalization. 10.1073/pnas

Duan, Y., Zhan, J., Gross, J., Ince, R. A. & Schyns, P. G. (2024). Pre-frontal cortex guides dimension-reducing transformations in the occipito-ventral pathway for categorization behaviors. Current Biology, 34, 3392–3404.

El-Gaby, M., Loyd Harris, A., R Whittington, J. C., Bhomick, A., Walton, M. E., Akam, T., & J Behrens, T. E. (2023). A Cellular Basis for Mapping Behavioural Structure. BioArxiv. 10.1101/2023.11.04.565609

Flesch, T., Juechems, K., Dumbalska, T., Saxe, A., & Summerfield, C. (2022). Orthogonal representations for robust context-dependent task performance in brains and neural networks. Neuron, 110(7), 1258–1270.e11. 10.1016/j.neuron.2022.01.005

Flesch, T., Saxe, A., & Summerfield, C. (2023). Continual task learning in natural and artificial agents. In Trends in Neurosciences (Vol. 46, Issue 3, pp. 199–210). Elsevier Ltd. 10.1016/j.tins.2022.12.006

Garner, K., Lynch, C. R. & Dux, P. E. (2016). Transfer of training benefits requires rules we cannot see (or hear). Journal of Experimental Psychology: Human Perception and Performance, 42, 1148.

Garvert, M. M., Dolan, R. J., & Behrens, T. E. (2017). A map of abstract relational knowledge in the human hippocampal–entorhinal cortex. eLife, 6, 1–20. 10.7554/elife.17086

Garvert, M. M., Saanum, T., Schulz, E., Schuck, N. W., & Doeller, C. F. (2023). Hippocampal spatio-predictive cognitive maps adaptively guide reward generalization. Nature Neuroscience, 26(4), 615–626. 10.1038/s41593-023-01283-x

Henin, S., Turk-Browne, N. B., Friedman, D., Liu, A., Dugan, P., Flinker, A., Doyle, W., Devinsky, O., & Melloni, L. (2021). Learning hierarchical sequence representations across human cortex and hippocampus. Science Advances, 7(8), 1–13. 10.1126/sciadv.abc4530

Holton, E., Braun, L., Thompson, J., Grohn, J. & Summerfield, C. (2026). Humans and neural networks show similar patterns of transfer and interference during continual learning. Nature Human Behaviour 10, 111–125. 10.1038/s41562-025-02318-y

Huys, Q. J. M., Maia, T. V., & Frank, M. J. (2016). Computational psychiatry as a bridge from neuroscience to clinical applications. In Nature Neuroscience (Vol. 19, Issue 3, pp. 404–413). Nature Publishing Group. 10.1038/nn.4238

Johnston, W. J., & Fusi, S. (2023). Abstract representations emerge naturally in neural networks trained to perform multiple tasks. Nature Communications, 14(1). 10.1038/s41467-023-36583-0

Kazanina, N., & Poeppel, D. (2023). The neural ingredients for a language of thought are available. In Trends in Cognitive Sciences (Vol. 27, Issue 11, pp. 996–1007). Elsevier Ltd. 10.1016/j.tics.2023.07.012

Kriegeskorte, N., Mur, M., & Bandettini, P. (2008). Representational similarity analysis – connecting the branches of systems neuroscience. Frontiers in Systems Neuroscience, 2(November), 1–28. 10.3389/neuro.06.004.2008

Kumar, S., Dasgupta, I., Marjieh, R., Daw, N. D., Cohen, J. D., & Griffiths, T. L. (2022). Disentangling Abstraction from Statistical Pattern Matching in Human and Machine Learning. ArXiv. http://arxiv.org/abs/2204.01437

Kurth-Nelson, Z., Behrens, T., Wayne, G., Miller, K., Luettgau, L., Dolan, R., Liu, Y., & Schwartenbeck, P. (2023). Replay and compositional computation. In Neuron (Vol. 111, Issue 4, pp. 454–469). Cell Press. 10.1016/j.neuron.2022.12.028

Lake, B. M., Salakhutdinov, R., & Tenenbaum, J. B. (2015). Human-level concept learning through probabilistic program induction. Science, 350(6266), 1332–1338. 10.1126/science.aab3050

Lake, B. M., Ullman, T. D., Tenenbaum, J. B., & Gershman, S. J. (2017). Building machines that learn and think like people. Behavioral and Brain Sciences, 40(2017). 10.1017/S0140525X16001837

Lehnert, L., Littman, M. L., & Frank, M. J. (2020). Reward-predictive representations generalize across tasks in reinforcement learning. PLoS Computational Biology, 16(10), 1–27. 10.1371/journal.pcbi.1008317

Luettgau, L., Erdmann, T., Veselic, S., Stachenfeld, K. L., Kurth-Nelson, Z., Moran, R., & Dolan, R. J. (2024). Decomposing dynamical subprocesses for compositional generalization. Proceedings of the National Academy of Sciences, 121(46). 10.1073/pnas.2408134121

Luettgau, L., Porcu, E., Tempelmann, C., & Jocham, G. (2022). Reinstatement of Cortical Outcome Representations during Higher-Order Learning. Cerebral Cortex, 32(1), 93–109. 10.1093/cercor/bhab196

Luettgau, L., Tempelmann, C., Kaiser, L. F., & Jocham, G. (2020). Decisions bias future choices by modifying hippocampal associative memories. Nature Communications, 11, 3318. 10.1101/802462

Luyckx, F., Nili, H., Spitzer, B. & Summerfield, C. (2019). Neural structure mapping in human probabilistic reward learning. eLife, 8, e42816. 10.7554/eLife.42816

Mark, S., Moran, R., Parr, T., Kennerley, S. W., & Behrens, T. E. J. (2020). Transferring structural knowledge across cognitive maps in humans and models. Nature Communications, 11(1), 1–12. 10.1038/s41467-020-18254-6

McElreath, R. (2020a). *rethinking: Statstical rethinking book package* (R package version 2.00).

McElreath, R. (2020b). Statistical Rethinking (2nd ed.). CRC Press. 10.1080/09332480.2017.1302722

Murphy, G. L. (1988). Comprehending Complex Concepts. Cognitive Science, 12(4), 529–562. 10.1207/s15516709cog1204_2

Nieh, E. H., Schottdorf, M., Freeman, N. W., Low, R. J., Lewallen, S., Koay, S. A., Pinto, L., Gauthier, J. L., Brody, C. D., & Tank, D. W. (2021). Geometry of abstract learned knowledge in the hippocampus. Nature, 595(7865), 80–84. 10.1038/s41586-021-03652-7

Nili, H., Wingfield, C., Walther, A., Su, L., Marslen-Wilson, W. & Kriegeskorte, N. (2014). A toolbox for representational similarity analysis. PLoS Computational Biology, 10, e1003553.10.1371/journal.pcbi.1003553

Nour, M. M., Liu, Y., Arumuham, A., Kurth-Nelson, Z., & Dolan, R. J. (2021). Impaired neural replay of inferred relationships in schizophrenia. Cell, 184(16), 4315–4328.e17. 10.1016/j.cell.2021.06.012

O’Donnell, T. J., Goodman, N. D., & Tenenbaum, J. B. (2009). Fragment Grammars: Exploring Computation and Reuse in Language. www.csail.mit.edu

O’Donnell, T. J., Snedeker, J., Tenenbaum, J. B., & Goodman, N. D. (2011). Productivity and Reuse in Language. Proceedings of Cognitive Science Conference.

Pesnot Lerousseau, J., & Summerfield, C. (2024). Space as a scaffold for rotational generalisation of abstract concepts. ELife, 13. 10.7554/eLife.93636

Reber, A. S. (1967). Implicit Learning of Artificial Grammars 1. In JOURNAL OF VERBAL LEARNING AND VERBAL BEHAVIOR (Vol. 6).

Redish, A. D. (2016). Vicarious trial and error. In Nature Reviews Neuroscience (Vol. 17, Issue 3, pp. 147–159). Nature Publishing Group. 10.1038/nrn.2015.30

Rmus, M., Ritz, H., Hunter, L. E., Bornstein, A. M., & Shenhav, A. (2022). Humans can navigate complex graph structures acquired during latent learning. Cognition, 225. 10.1016/j.cognition.2022.105103

RStudioTeam. (2019). RStudio: Integrated Development for R (1.2.5033). RStudio, Inc. http://www.rstudio.com/

Rubino, V., Hamidi, M., Dayan, P., & Wu, C. M. (2023). Compositionality under time pressure. Proceedings of the Annual Meeting of the Cognitive Science Society, 45.

Schapiro, A. C., Rogers, T. T., Cordova, N. I., Turk-Browne, N. B., & Botvinick, M. M. (2013). Neural representations of events arise from temporal community structure. Nature Neuroscience, 16(4), 486–492. 10.1038/nn.3331

Schwartenbeck, P., Baram, A., Liu, Y., Mark, S., Muller, T., Dolan, R., Botvinick, M., Kurth-Nelson, Z., & Behrens, T. (2023). Generative replay underlies compositional inference in the hippocampal-prefrontal circuit. Cell, 186(22), 4885–4897.e14. 10.1016/j.cell.2023.09.004

Sheahan, H., Luyckx, F., Nelli, S., Teupe, C., & Summerfield, C. (2021). Neural state space alignment for magnitude generalization in humans and recurrent networks. Neuron, 109(7), 1214–1226.e8. 10.1016/j.neuron.2021.02.004

Shepard, R. N. (1987). Toward a Universal Law of Generalization for Psychological Science. Science, 237(4820). 10.1126/science.3629243

Sherman, B. E., Graves, K. N., & Turk-Browne, N. B. (2020). The prevalence and importance of statistical learning in human cognition and behavior. Current Opinion in Behavioral Sciences, 32, 15–20. 10.1016/j.cobeha.2020.01.015

Sorscher, B., Ganguli, S., & Sompolinsky, H. (2022). Neural representational geometry underlies few-shot concept learning. Proceedings of the National Academy of Sciences. 10.1073/pnas

Stan Development Team. (2020). *RStan: the R interface to Stan* (R package version 2.19.3.). http://mc-stan.org/

Story, G. W., Smith, R., Moutoussis, M., Isabel M. Berwian, I. M., Nolte, T., Bilek, E., Jenifer Z. Siegel, J. Z., & Dolan, R. J. (2024). A Social Inference Model of Idealization and Devaluation. Psychological Review, 131(3), 749–780. 10.1037/rev0000430

Teichmann, L., Grootswagers, T., Carlson, T. & Rich, A. N. (2018). Decoding digits and dice with magnetoencephalography: evidence for a shared representation of magnitude. Journal of Cognitive Neuroscience, 30(7), 999–1010

Tsividis, P. A., Loula, J., Burga, J., Foss, N., Campero, A., Pouncy, T., Gershman, S. J., & Tenenbaum, J. B. (2021). Human-Level Reinforcement Learning through Theory-Based Modeling, Exploration, and Planning. ArXiv, 1–67. http://arxiv.org/abs/2107.12544

Turk-Browne, N. B., Isola, P. J., Scholl, B. J., & Treat, T. A. (2008). Multidimensional Visual Statistical Learning. Journal of Experimental Psychology: Learning Memory and Cognition, 34(2), 399–407. 10.1037/0278-7393.34.2.399

Whittington, J. C. R., McCaffary, D., Bakermans, J. J. W., & Behrens, T. E. J. (2022). How to build a cognitive map: insights from models of the hippocampal formation. 1–25. http://arxiv.org/abs/2202.01682

Whittington, J. C. R., Muller, T. H., Mark, S., Chen, G., Barry, C., Burgess, N., & Behrens, T. E. J. (2020). The Tolman-Eichenbaum Machine: Unifying Space and Relational Memory through Generalization in the Hippocampal Formation. Cell, 183(5), 1249–1263.e23. 10.1016/j.cell.2020.10.024

Yang, G. R., Joglekar, M. R., Song, H. F., Newsome, W. T., & Wang, X. J. (2019). Task representations in neural networks trained to perform many cognitive tasks. Nature Neuroscience, 22(2), 297–306. 10.1038/s41593-018-0310-2

Zhang, M. & Yu, Q. (2024). The representation of abstract goals in working memory is supported by task-congruent neural geometry. PLoS Biology, 22, e3002461.

Zheng, X. Y., Hebart, M. N., Grill, F., Dolan, R. J., Doeller, C. F., Cools, R., & Garvert, M. M. (2024). Parallel cognitive maps for multiple knowledge structures in the hippocampal formation. Cerebral Cortex, 34(2). 10.1093/cercor/bhad485

Zhou, J., Jia, C., Montesinos-Cartagena, M., Gardner, M. P. H., Zong, W., & Schoenbaum, G. (2021). Evolving schema representations in orbitofrontal ensembles during learning. Nature, 590(7847), 606–611. 10.1038/s41586-020-03061-2

## References

Huang, Q., & Luo, H. (2024). Shared structure facilitates working memory of multiple sequences. ELife, 12. 10.7554/eLife.93158

Kern, S., Nagel, J., Gerchen, M. F., Guersoy, C., Meyer-Lin-Denberg, A., Kirsch, P., Dolan, R. J., Gais, S., & Feld, G. B. (2024). Reactivation strength during cued recall is modulated by graph distance within cognitive maps. ELife. 10.7554/eLife.93357.3

Kurth-Nelson, Z., Economides, M., Dolan, R. J., & Dayan, P. (2016). Fast Sequences of Non-spatial State Representations in Humans. Neuron, 91(1), 194–204. 10.1016/j.neuron.2016.05.028

Liu, Y., Dolan, R. J., Kurth-Nelson, Z., & Behrens, T. E. J. (2019). Human Replay Spontaneously Reorganizes Experience. Cell, 178, 1–13. 10.1016/j.cell.2019.06.012

Liu, Y., Mattar, M. G., Behrens, T. E. J., Daw, N. D., & Dolan, R. J. (2021). Experience replay is associated with efficient nonlocal learning. Science, 372(6544). 10.1126/science.abf1357

